# Chromosome-level genome and caste transcriptomes for Africa’s most widespread social wasp (*Belonogaster juncea*) reveal unexpected genomic diversity

**DOI:** 10.64898/2026.10.16.759595

**Authors:** Christopher D.R. Wyatt, Emeline Favreau, Katie Geist, Rick Masonbrink, Fernando Duarte, Cintia Akemi Oi, Benjamin Aaron Taylor, Romuald Tcheutchoua, Sarielle Kapmegne, Alex Cerqueira De Araujo, Maurice Tindo, Paul Serge Masse, Patrick Kennedy, Amy Toth, Sandra Rehan, Seirian Sumner

**Affiliations:** Centre for Biodiversity and Environment Research, Department of Genetics, Evolution and Environment, University College London, London, UK; Department of Medicine, University of Cambridge, Jeffrey Cheah Biomedical Centre, Cambridge Biomedical Campus, Cambridge, CB2 0AW, UK; Department of Ecology, Evolution and Organismal Biology, and Department of Plant Pathology, Entomology, and Microbiology, Iowa State University, Ames, IA 50011, USA; Genome Informatics Facility, Iowa State University, 449 Bessey Hall, Osborne Rd, Ames, IA 50011, USA; Department of Animal Biology and Physiology, Faculty of Science, University of Douala, BP 24157, Douala, Cameroon; Department of Animal Biology and Physiology, Faculty of Science, University of Yaoundé 1, BP 812, Yaoundé, Cameroon; School of Biological Sciences, University of Bristol, 24 Tyndall Avenue, Bristol, BS8 1TQ, UK; Department of Biology, York University, Toronto, ON, M3J 1P3, Canada

**Keywords:** *Belonogaster juncea*, genome assembly, social wasp, eusociality, caste differentiation, RNA-seq

## Abstract

*Belonogaster* is an African genus of primitively eusocial wasps. They are widespread and conspicuous across sub-Saharan Africa and yet are among the least studied groups of social wasps. Here, we present the first high-quality, chromosome-level genome assembly for a representative of this genus, *Belonogaster juncea*, complemented by an RNA-seq analysis of brain transcription among individuals representing reproductive and non-reproductive castes. The assembled genome spans 315.8 Mb with a scaffold N50 of 7.91 Mb. The annotation contained 15,159 protein-coding genes and 20,046 isoforms, with Benchmarking Universal Single-Copy Orthologs (BUSCO) completeness of 98.0% for the genome and 97.8% for the annotation. Using this new assembly, we show that the *B. juncea* genome differs notably from other social wasp genomes: it has the highest putative chromosome number in a sequenced wasp genome to date and it shows high levels of syntenic breaks. We find caste neuro-transcription differences between queens and workers, revealing 573 differentially expressed genes (341 worker biased, 232 queen biased), with significant overlap in caste-specific genes when compared to *Polistes dominula* and *Liostenogaster flavolineata*. This genomic resource provides a foundation for understanding the molecular mechanisms underlying primitive eusociality and social evolution in wasps, and opens up new research opportunities for a genus of little-studied insects from the Global South.

---

Social wasps are classic models for understanding the evolution of cooperation and conflict^1,2^. With over 1,000 species globally, social wasps span a vast diversity of social structures, levels of social complexity, and cooperative behaviours^3^. However, our understanding of wasp social evolution is constrained by the sparse and unbalanced availability of high-quality genome sequences. Across the polistine wasps, there are currently only sequenced genomes available for eight species^4–8^, of which seven are within the intensely studied genus *Polistes.* More generally across insects, the availability of genomic resources remains highly biased towards the Global North, leaving the insect biodiversity of the Global South – especially African species - underrepresented^9^. Here, to extend genomic resources for the study of social evolution, we provide a chromosome-level genome and transcriptome analysis of the most abundant and widespread social wasp of sub-Saharan Africa.

The needle-waist wasp *Belonogaster juncea* (*B. juncea*; Vespidae: Polistinae; Fig. 1a) is among Africa’s most familiar insects^10–13^. The species has a wide and climatically cosmopolitan distribution over a range of around 40 degrees latitude^14^, encompassing lowland tropical rainforests, savannas and temperate highlands. As a simple eusocial insect, *B. juncea* offers a model for understanding the origin and elaboration of eusociality. Despite early claims that *B. juncea* comprised groups of multiple egg-laying females^15^ (polygyny), *Belonogaster* colonies exhibit a similar social structure to *Polistes* wasps^16–18,11,19^. Typically, a single egg-laying female (the queen) is helped to establish a nest by subordinate females^13^ (co-foundresses). Many of her female offspring remain on the natal nest as workers, helping rear their siblings. Colony sizes are small, with most mature nests rarely rising above 30 females. The queen maintains her reproductive monopoly by presiding over a dominance hierarchy within the group^17,11^.

**Figure 1.**
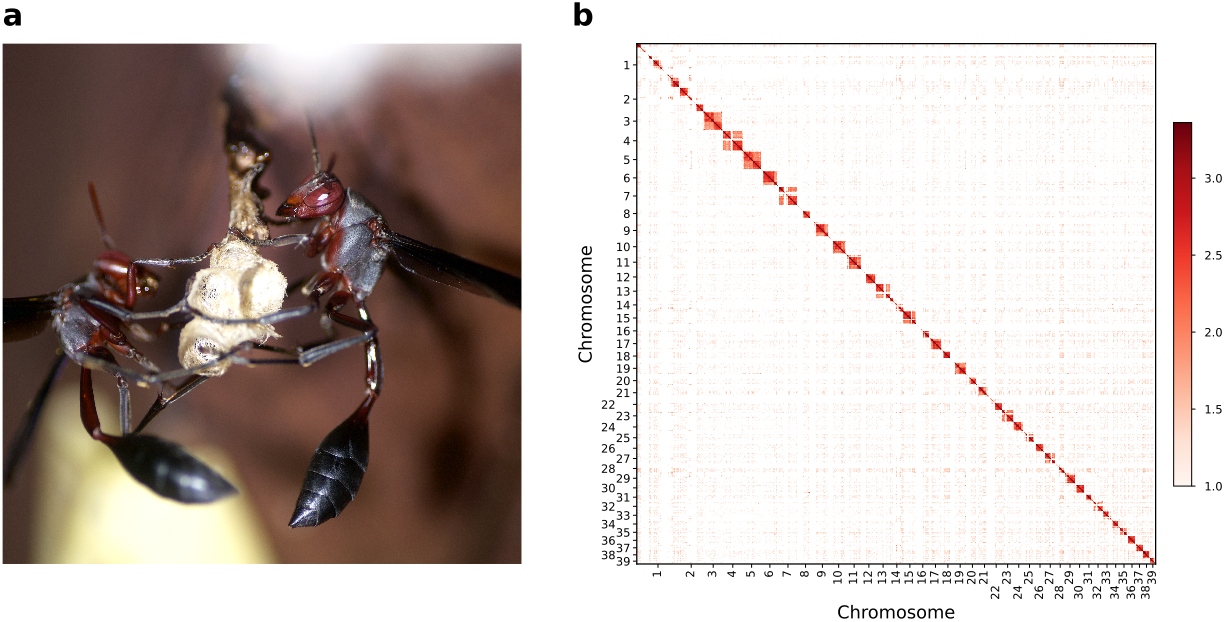
Genomic resources for Africa’s most familiar social wasps. (a) *Belonogaster* wasps are abundant across Africa. Nests are small and primitively eusocial (photograph: Patrick Kennedy, Yaoundé, Cameroon). (b) Omni-C contact map of the *B. juncea* chromosome-level assembly, showing the 39 chromosome-scale scaffolds (HiC_scaffold_1–39) in numerical order (311.3 Mb, 98.6% of the assembly; 60 small unplaced scaffolds, 4.5 Mb, are not shown). Colour shows the number of Hi-C contacts between 250-kb bins (log₁₀ scale); bin pairs with fewer than 10 contacts (the 75th percentile of between-chromosome contacts) are white.

*B. juncea* spans a remarkable range of climatic zones, and its life history varies accordingly across the continent. Two intergrading subspecies are currently recognised^14^, differing in wing colouration: the more-tropical *B. j. juncea* (with dark and iridescent wings) and the more-temperate *B. j. colonialis* (with transparent and brown-tipped wings). In the Congo Basin, sustained rainfall allows *B. j. juncea* to construct nests continuously throughout the year^12^. In temperate southern Africa, by contrast, *Belonogaster* wasps (including *B. j. colonialis*) follow a strictly seasonal nest cycle^16,18^. Here, young females emerge from a four-month hibernation to establish new nests as queens in the spring, and nests fail with the onset of winter. At present, little is known of *Belonogaster* behaviour and natural history in Africa’s arid and semi-arid zones, including the Sahel and East African savannas.

Unlike the extensively studied genus *Polistes*^2^*, Belonogaster* has received sporadic attention from evolutionary biologists. The first dedicated taxonomy was by du Buysson in 1910^20^. Six years later, the genus enjoyed brief celebrity in the nascent field of social evolution, when Emile Roubaud published behavioural observations of colonies in Dahomey (modern-day Benin)^15^. Roubaud’s discovery of larval-adult trophallaxis (larvae feeding adults by regurgitating their saliva) inspired his influential^21^ – now discarded – early theory^15,22^ that wasps evolved sociality due to adults becoming addicted to a life-giving larval ‘elixir’^23^. Over the subsequent century, however, Africa’s most widespread wasps were largely neglected in comparison to other genera, with three notable and valuable exceptions. In Somalia, Pardi and Marino Piccioli examined the nest architecture^24^ and behaviour^25^ of *B. grisea*, suggesting that queens and workers may differ not only in behaviour but also in morphometrics^26,27^ (pre-imaginal caste biasing). In South Africa, Keeping examined the social behaviour of *B. petiolata*, documenting similar caste differentiation^28^ and the seasonal nesting cycle typical of *Belonogaster* in temperate latitudes^16–18,28^; more recently, Kelstrup and coauthors^19^ quantified hormone titres in the South African species *B. longitarsus.* In Cameroon, Tindo and coauthors studied the social behaviour of *B. juncea*, revealing that subordinate helpers boost nest success^13^, that queens maintain dominance by aggression^11^, and that dead queens can be replaced by subordinate nest-mates in long-lived colonies^12^. The last major taxonomic revision of the genus was in 1982^14^. Although much remains unknown, comparative phylogenetic studies^29^ of sociality across wasps have used *Belonogaster* to reconstruct ancestral states. *Belonogaster* offers an evolutionarily significant taxon for comparative genomics with other social wasps, constrained to date by an absence of genomic or genetic resources.

Here, we provide a high-quality chromosome-level genome assembly for the species *B. juncea* (subspecies *juncea*), and characterise gene content and repetitive elements. Alongside our genome assembly, we explore chromosomal functional enrichment, gene family evolution and syntenic relationships with closely related species. We also provide brain transcriptomes for queens and workers, and compare patterns of caste-specific gene expression with reproductive and non-reproductive castes of other similarly simple social wasp species. Our results expand the current landscape for comparative genomics in social wasps to a long-neglected but evolutionarily significant taxon in the Global South.

## Materials and Methods

### Genome extraction and library preparation

An adult female of *B. juncea* was collected from a post-emergence nest on 8th April 2021 in Soa, near Yaoundé (Cameroon) into 95% ethanol and sequenced (Cantata Bio). Qiagen kit for PacBio DNA extraction was used for the extraction using manufacturer protocol. The SMRTbell Express Template Prep kit 2.0 (PacBio, Menlo Park, CA, USA) was employed to prepare the PacBio sequencing library in accordance with the manufacturer’s protocol. The library was placed onto a PacBio Sequel II sequencer with 8 M SMRT cell after being bound to polymerase using the Sequel II Binding Kit 2.0 (PacBio). For each Omni-C (proximity-ligation Hi-C) library, chromatin was fixed in place with formaldehyde within the nucleus and then digested with DNase I. Chromatin ends were repaired and ligated to a biotinylated bridge adapter. The adapter-containing ends were then proximity ligated. The DNA was purified and crosslinks were reversed by proximity ligation. The purification was treated to remove biotin that was not intrinsic to the ligate segments. The kit from New England Biolabs Next Ultra enzymes and Illumina-compatible adapters was used for sequencing the libraries. Biotinylated fragments were captured on streptavidin beads prior to PCR enrichment, and the resulting Omni-C library was sequenced on an Illumina HiSeq X platform.

### Genome assembly

Cantata Bio (Formerly Dovetail Genomics) constructed the genome assembly using Omni-C data and PacBio HiFi reads. Primary contig assembly was performed using hifiasm^30^ as part of Cantata Bio Genomics’ proprietary assembly pipeline, followed by scaffolding with HiRise using Omni-C read pairs. Omni-C read pairs with mapping quality >50 were used in conjunction with PacBio HiFi reads for de novo assembly using wtdbg2^31^ v2.5. PacBio HIFI reads were checked for contamination using BlobTools^32^ v1.1.1 (within the Cantata Bio report: https://doi.org/10.5281/zenodo.18259073). Read length distributions were visualised using NanoPack^33^.

Omni-C reads were aligned to the draft assembly using BWA^34^, and HiRise was used to model genomic distance between read pairs, identify and break putative misjoins, and make scaffold joins exceeding confidence thresholds. For downstream Hi-C quality control and visualisation, Hi-C reads were extracted from BAM files using Picard^35^ v2.17.0, which were then processed with Juicer^36^ v1.5.7, 3D-DNA^37^ (version 180114), and manual curation (merging of scaffolds) using JuiceBox^38^ v1.11 after being aligned to scaffolds with BWA. The contact map in Figure 1b was drawn from the resulting Juicer contact map (.hic) and the curated assembly layout: contacts were extracted at 50-kb resolution with hic-straw v1.3.1, placed in the order and orientation of the final assembly, summed into 250-kb bins and plotted for the 39 chromosome-scale scaffolds (code: https://github.com/Sumner-lab/Belonogaster_genome_paper/tree/main/HiC_and_satellite).

The final genome assembly has been deposited in NCBI under accession PRJNA1357342 (BioSample: SAMN53082503; B_junceaGenome.FINAL.fasta.gz). Raw PacBio HiFi reads (run accession SRR39278268) and Omni-C reads (run accession SRR41179205) are available from the NCBI Sequence Read Archive under BioProject PRJNA1357342^39^.

### Genome assembly quality assessment

nf-core/genomeqc^40^ (v1.0.1) was used to assess the quality of the *B. juncea* final assembly and 11 other Vespidae assemblies (Supplementary table 1: Comparison genomes): eight species from the subfamily Vespinae, one from Polistinae, one from Eumeninae, and one from Stenogastrinae. Genomes and annotations were obtained from RefSeq and the Darwin Tree of Life project, together with three Hi-C assemblies from the Sumner, Toth and Rehan labs: *L. flavolineata*^41^, *P. dominula*^42^ and *Vespa crabro*^43^. General assembly statistics were computed with QUAST^44^ v5.3.0, and completeness was assessed with BUSCO^45^ v6.1.0 against the hymenoptera_odb10 lineage dataset. Transposable elements were annotated in all 12 assemblies with RepeatMasker^46^ v4.1.5 (rush mode, -qq) using the Dfam^47^ v3.9 families for Hymenoptera (curated and un-curated, including species-specific libraries for 10 of the 12 species but not for *B. juncea* or *L. flavolineata*; extracted with FamDB from Dfam partitions 0 and 3 and clustered with MMseqs2^48^ easy-linclust). No de novo repeat library was added, so that all species were compared against the same set of known families (Figure 2; Supplementary Table 1: RepeatMasker). To test whether *B. juncea* gene models were derived from transposable elements, we calculated the proportion of each gene’s coding sequence that overlapped interspersed repeats annotated by RepeatMasker^46^.

**Figure 2.**
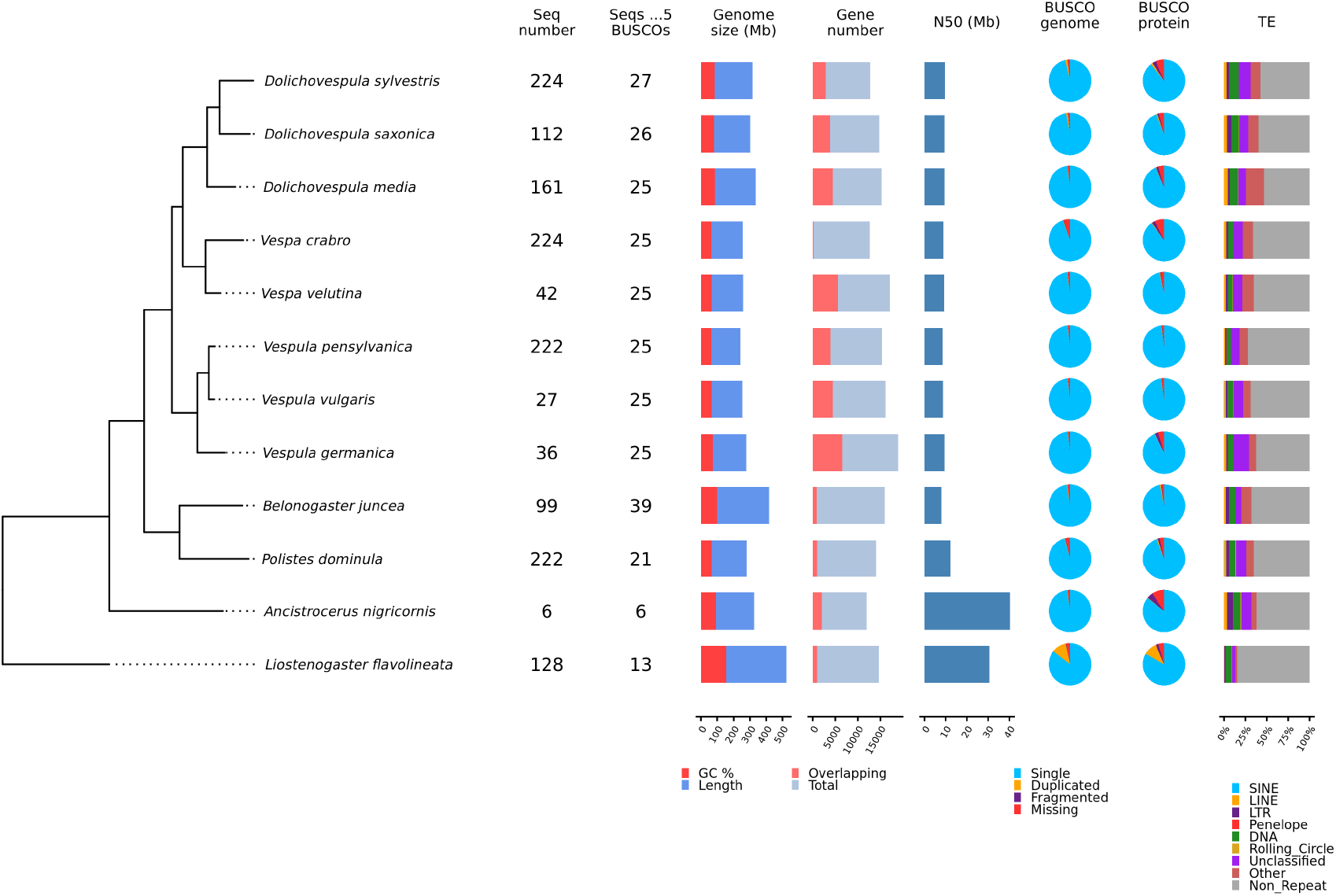
Phylogeny and genome statistics of the 12 vespid assemblies. Species tree inferred by OrthoFinder from the predicted proteins (left), with summary statistics from nf-core/genomeqc (right). “Seq number”, number of scaffolds or chromosomes in the assembly; “Seqs >5 BUSCOs”, number of scaffolds containing more than five BUSCO genes, an indication of chromosome number. “Genome size”, assembly length, with the GC fraction in red. “Gene number”, number of genes, with the number overlapping another gene (on either strand) in red. “N50”, scaffold N50. BUSCO pie charts show completeness of the genome (“genome”) and of the predicted proteins (“protein”) against hymenoptera_odb10: single-copy (light blue), duplicated (orange), fragmented (purple) and missing (red). “TE”, proportion of each genome masked by RepeatMasker with the Dfam families for Hymenoptera, by repeat class (“Other”: simple repeats, low-complexity regions, satellites and small RNAs; grey, not masked).

### Tandem repeats and satellite DNA

Tandem repeats in the *B. juncea* assembly were identified with Tandem Repeats Finder^49^ v4.09.1 (parameters 2 7 7 80 10 50 2000, maximum array length 10 Mb). Among them were many arrays of a satellite with a 296-bp repeat unit; arrays with repeat units of 285– 305 bp, or two or three times that length, were counted as this satellite (consensus sequence in the code repository, below). The *B. juncea* assembly has 685 such arrays, on all 39 chromosomes. To test whether other wasps carry the satellite, we searched the other 11 vespid assemblies for its sequence, counting exact matches to its 12-mers (both strands) in 2-kb windows. None of them contains it: no window in any of these assemblies had more than 2% of its positions matching. We also compared where the satellite arrays lie with the Omni-C contact map, in 50-kb windows. Code: https://github.com/Sumner-lab/Belonogaster_genome_paper/tree/main/HiC_and_satellite.

### Genome annotation

Two versions of the gene annotation were generated for *B. juncea*, *P. dominula* and *L. flavolineata*, and both are available on Zenodo with the genome assemblies and protein sequences (*B. juncea*: https://doi.org/10.5281/zenodo.18892456; *P. dominula*: https://doi.org/10.5281/zenodo.23264930; *L. flavolineata*: https://doi.org/10.5281/zenodo.23266517). All analyses in this study use V2 (files *.V2.gff). V1 is retained because it has been used by third-party publications; V2 is more complete, using the latest versions of the tools.

Annotation V1. Repeats were identified with RepeatModeler^50^ v2.0.2 and soft-masked with RepeatMasker^46^ v4.1.2-p1. RNA extracted from the heads, thoraxes and abdomens of three females (samples “Btho1”, “Btho2”, “Btho3”, “Babd2”, “Bhead1”, “Bhead2”, “Bhead3”; GSE320383) was aligned to the genome using STAR^51^ v2.7.6a. AUGUSTUS^52^ v3.4 was trained with the BUSCO v5.12 hymenoptera_odb10 lineage dataset, using RNA-seq-derived hints for CDS, exons and introns. Gene models were then generated with BRAKER^53^ v3.0.2 and GeneMark^54^ v4.69 and refined using GenomeThreader^55^ alignments of proteins from NCBI Annotation Release 103. Predicted proteins were functionally annotated by alignment against the NCBI nr database with DIAMOND^56^ v2.0.25. The V1 annotation is available on Zenodo (https://doi.org/10.5281/zenodo.18892456; file rd014j_Belonogaster_juncea.V1.gff).

Annotation V2. Repeats were annotated with HiTE^57^ v3.1.2 using curated metazoan sequences from Dfam^47^ v3.8 and masked with BEDTools^58^ v2.31.1. The same RNA-seq reads were trimmed with Trimmomatic^59^ v0.39 (parameters: -phred33 ILLUMINACLIP:TruSeq3-PE.fa:2:30:10 LEADING:3 TRAILING:3 SLIDINGWINDOW:4:15 MINLEN:36), quality-checked with FastQC v0.12.1 and MultiQC^60^ v1.25.2, mapped to the genome with hisat2^61^ v2.2.1, and sorted with samtools^62^ v1.21. Gene prediction was performed by AUGUSTUS^63^ v3.5.0, BRAKER3^64^ v3.0.8 and GeneMark-ETP^65^ v1.02, using a protein database of sequences extracted from hymenopteran RefSeq. The pipeline was run in ETP mode. TSEBRA^66^ was configured as follows: P 5; E 10; C 10; M 5; intron_support 0.2; stasto_support 1; e_1 0.3; e_2 1; e_3 0.2; e_4 0.2; e_5 0.18; e_6 0.18. The best_by_compleasm.py v1.0.1 script (from BRAKER) was then used to keep non-spurious models filtered out by TSEBRA. Nucleotide and amino acid CDS sequences were then extracted using AGAT^67^ v1.4.1. *P. dominula* and *L. flavolineata* were annotated using the same tools, as listed in their genome notes^41,42^. For all three species, the V2 annotation (*.V2.gff in the Zenodo records above) was used throughout this manuscript.

### Genome annotation quality assessment

Annotation quality and completeness of both *B. juncea* and the other 11 Vespidae species were assessed using nf-core/genomeqc^40^ (v1.0.1). General annotation statistics were obtained using AGAT^67^ v1.6.1. For completeness, BUSCO^45^ v6.1.0 and the hymenoptera_odb10 lineage database were used. To evaluate completeness, the longest isoform was selected for every gene and GFFREAD^68^ v0.12.7 was used to extract coding sequences from the assembly. Finally, using the extracted sequences as input, BUSCO was run on protein mode. OrthoFinder^69^ v2.5.5 was used to perform phylogenetic orthology inference on the extracted coding sequences. OrthoFinder identified orthogroups for all species, inferred gene trees for these orthogroups, and identified the species tree from these gene trees.

### Mitochondrial genome

To assemble the annotated mitochondrial genome, we used MitoHiFi^70^ v2 in reads mode on a randomly-selected subset of the raw reads, which is suited to PacBio HiFi sequences. We used MitoHiFi’s default annotator, MitoFinder^71^. To supply a related reference to MitoHiFi to guide read mapping, we used the *Polistes rothneyi* complete mitochondrion (NCBI accession: NC_063693.1). Mean depth of the mapped reads used for the assembly (calculated with minimap2 v2.30) was 655.45.

### RNA-seq analysis

*B. juncea* samples for transcriptome sequencing were collected from Yaoundé (n=15) and Soa (n=10) (Cameroon) in 2021. The nests were collected and individual wasps were placed in Eppendorf tubes containing RNAlater and stored at -20C. To assess reproductive state, the ovaries for each female were dissected under a stereomicroscope (Leica): females with developed oocytes were classed as reproductives; those without developed oocytes were classed as non-reproductives. Additionally, caste was verified using behavioural observation, using egg-laying and residency on nest to determine the queen. In total, 14 females were classed as reproductives (10 from Soa and 4 from Yaoundé) and 11 as non-reproductives (all from Yaoundé).

Wasp brains were dissected out of their head capsules in RNALater fluid, and RNA from 25 brains were extracted using the Qiagen RNAsy Plus Universal Mini Kit according to the manufacturer’s protocol. The samples were quantified using NanoDrop.

The samples of *B. juncea* were sequenced (RNA-seq, Novogene), using the NEB Next UltraTM RNA Library Prep kit on a NovaSeq 6000 machine. Raw reads were deposited NCBI PRJNA1427691;GSE320383 (Sample details in Supplementary Table 2: *Belonogaster* sample details).

We mapped transcripts and quantified gene expression in the brains of reproductive and nonreproductive *B. juncea* and compared these with equivalent published data available for two other simple social species: *Liostenogaster flavolineata (L. flavolineata)*^72^ and *Polistes dominula (P. dominula)*^73^ (whole brains from the head capsules of individual adult females (including the suboesophageal ganglion) using the nf-core/rnaseq^74^ pipeline v3.21.0 in Nextflow^75^ v25.10.0 with STAR^51^ and Salmon^76^. Quality of the mapping and quantification was assessed using multiQC^60^. Salmon counts for the adult female brain samples are deposited on GEO: GSE320383. MultiQC summaries for each species are available in Zenodo: https://doi.org/10.5281/zenodo.20629097).

We analysed differential expression in *R* using DESeq2^77^ v1.42.1. Prior to this analysis, we filtered out genes with low expression, independently for each species: first, we removed genes whose mean expression across all samples belonging to that species was lower than 1 read/sample; second, we removed genes whose mean expression was below 5 reads/sample within both groups of individuals (reproductives and nonreproductives). We then identified differentially expressed genes (DEGs) between reproductives and nonreproductives using a local model fit. For principal component analysis, counts were variance-stabilised (DESeq2 vst, blind to caste) and the 500 most variable genes were used. We considered that genes were differentially expressed in a given comparison if p < 0.05 after false discovery rate correction (Benjamini and Hochberg 1995)^78^.

To compare caste-biased expression across species, we used the orthogroups inferred by OrthoFinder^69^ v3.1.4 in the Excon^79^ run used for the gene family analysis (12 species; 11,702 orthogroups; see Gene family expansion and contraction). For each species and caste, we listed the orthogroups containing at least one gene upregulated in that caste. For each pair of species, we tested whether the same caste shared more orthogroups than expected by chance with a one-sided hypergeometric test, using as the universe the orthogroups that contained at least one gene tested for differential expression in both species (7,847–8,238 orthogroups). *P*-values were corrected across the six same-caste comparisons with the Benjamini–Hochberg method. Comparisons between opposite castes (queen-biased in one species and worker-biased in the other) served as a control. Code: https://github.com/Sumner-lab/Belonogaster_genome_paper.

### Functional annotation and GO enrichment

To determine the Gene Ontology (GO) enrichment of genes used in this genome, we annotated the genes using a Nextflow pipeline (Eco-flow/excon v2.5.0^79^), which creates protein sequences from each given genome and annotation and uses eggNOG-mapper v2.1.13^80^ to assign GO terms through orthology assignment (DIAMOND mode; taxonomic scope Insecta; bit score ≥ 60, ≥ 40% identity, ≥ 40% query and subject coverage). The GO files used in the genome-wide analyses are deposited in Zenodo (10.5281/zenodo.23416281). For the analysis of caste-biased genes, GO terms were assigned (excon v2.5.0, eggNOG-mapper v2.1.13) to the gene models used for RNA-seq quantification: 5,724 genes in *B. juncea* (431,497 annotations), 6,179 in *L. flavolineata* (470,584) and 5,493 in *P. dominula* (418,433) (Zenodo: 10.5281/zenodo.23174866).

To measure if specific gene sets were enriched for particular GO terms, we used the database of terms produced by eggNOG-mapper, then ran custom wrapper code (ChopGO, https://github.com/Sumner-lab/Belonogaster_genome_paper/blob/main/bin/ChopGO.pl), which runs TopGO. All P-values enriched are shown using Bonferroni correction and are considered significant if they were less than 0.05.

For the caste-biased gene sets, we tested enrichment directly in topGO v2.56.0 rather than with ChopGO, so that all tested GO terms were retained for multiple-testing correction (code:https://github.com/Sumner-lab/Belonogaster_genome_paper/tree/main/GeneOntologyOverlap). For each species, genes upregulated in reproductives (queen-biased) and in non-reproductives (worker-biased) were tested separately in each GO domain with one-sided Fisher’s exact tests (’classic’ algorithm). The background was the set of expressed genes (those retained after expression filtering) with at least one GO annotation (*B. juncea* 5,417; *L. flavolineata* 5,913; *P. dominula* 5,222 genes); using all annotated genes as the background gave similar results. P-values were corrected across all terms tested in each domain using the Benjamini–Hochberg method, and terms were considered significant at FDR < 0.05.

To test whether GO terms were enriched in two gene sets overlapped more than expected by chance (using a cut off of P < 0.01), we used a permutation test that accounts for the non-independence of GO terms. In each of 10,000 permutations, we drew random gene sets of the observed sizes from each species’ background (queen- and worker-biased sets without overlap), recomputed Fisher’s exact P-values for all GO terms, and took the same number of top-ranked terms as observed for each gene set. P-values were the proportion of permutations with at least the observed number of shared terms, adjusted across all pairwise comparisons with the Benjamini–Hochberg method.

### Chromosome GO enrichment

We tested each chromosome for enrichment of GO terms with the “chromosome_enrichment” function of Eco-flow/excon^79^ v2.5.1 (commit b809d7c), in the same run as the gene family analysis below. For each chromosome (scaffold) in each of the 12 species, the genes on that chromosome were compared with all GO-annotated genes in the genome (using topGO^81^ v2.56.0, classic algorithm, one-sided Fisher’s exact test, run separately for biological process, molecular function and cellular component). *P*-values were Bonferroni-corrected across all terms tested in each domain, and a term was considered enriched when its corrected *P* < 0.05 and at least 10 genes in the genome were annotated to it. Because clusters of tandemly duplicated genes can produce enrichment on their own, the test was repeated with duplicates collapsed, counting each orthogroup once per chromosome using the orthogroups from the same run. Full results are on Zenodo (https://doi.org/10.5281/zenodo.23221104).

### Gene family expansion and contraction

Gene family evolution was analysed in the same Excon^79^ v2.5.1 run (commit b809d7c) as the chromosome analyses. OrthoFinder v3.1.4 was run on the longest protein isoform of each gene in the 12 species to define orthogroups and infer a rooted species tree, whose branch lengths were rescaled and made ultrametric (ape chronoMPL) for CAFE5^82^. Orthogroups present in only one species or with 100 or more copies in any species were excluded, and 9,722 gene families were modelled. CAFE5 was run with an error model and with one to six gamma rate categories; four categories were selected by AIC, and a Poisson distribution of root family sizes was preferred over a uniform distribution by likelihood. A family was considered significantly expanded or contracted on a branch when its CAFE5 branch P ≤ 0.05. Because *L. flavolineata* is the only species on one side of the root, changes on its branch and on the branch leading to the other 11 species cannot be separated and depend on the model, so these two branches were not interpreted. To test robustness to the definition of gene families, the analysis was repeated with hierarchical orthogroups (N0) from OrthoFinder^69^ v2.5.5, keeping families absent at the root (11,781 families; three gamma categories and a Poisson root distribution selected as above). GO enrichment of the families expanded or contracted in each species (any change, no P-value cut-off) was tested against all orthogroups containing genes from that species with topGO (classic Fisher’s exact test), using GO terms assigned to orthogroups from the eggNOG-mapper annotations of their genes; P-values were corrected across all terms tested in each GO domain with the Benjamini–Hochberg method.

## Results and Discussion

### 1. Genome of Belonogaster juncea

#### Genome assembly and annotation

We generated a chromosome-scale assembly of the *B. juncea* (Figure 1a) genome using PacBio HiFi sequencing combined with Cantata Bio Omni-C proximity ligation. The PacBio library yielded 18.8 Gb of sequence data from 1,729,268 Circular Consensus Sequencing (CCS) reads, representing approximately 64-fold coverage based on an estimated genome size of 290 Mb. The Omni-C library generated approximately 30-fold sequence coverage from 19,174,176 high-quality read pairs (mapping quality >50) used for scaffolding.

Initial assembly with Hifiasm v0.15.4 produced a draft assembly of 410.7 Mb. Following contamination filtering with BlobTools v1.1.1 (Cantata Bio report: https://doi.org/10.5281/zenodo.18259073) and haplotig removal with purge_dups v1.2.5, the primary filtered assembly was reduced to 315.7 Mb organized into 236 scaffolds with a contig N50 of 3.85 Mb, L50 of 32, and N90 of 0.79 Mb. Further, Hi-C scaffolding with Cantata Bio HiRise using Omni-C proximity ligation data produced a chromosome-scale assembly of 315.7 Mb organized into 206 scaffolds with 46 gaps, yielding an N50 of 4.93 Mb, L50 of 21, and N90 of 0.84 Mb (HiRise report: https://doi.org/10.5281/zenodo.18259073). HiRise made 31 joins and 1 break to the input assembly. The largest scaffold in this assembly was 11.5 Mb, representing a 1.6-fold improvement over the input assembly (7.3 Mb).

Additional merging of scaffolds was performed using Juicebox, which reduced the numbers of scaffolds down to a final count of 99. Our final genome version of the *B. juncea* assembly had a scaffold N50 of 7.9 Mb, an L50 of 14, and an N90 of 5.36 Mb with a total length of 315.8 Mb (size difference likely due to addition of gaps after HiC super scaffolding) and a GC content was 32.05 % (**Supplementary Table 1:** Quast). Genome completeness assessed against the hymenoptera_odb10 BUSCO database (5,991 genes) was 98.0% (BUSCO v6.1.0; Supplementary Table 1: BUSCO genome). Taken together, the assembly statistics demonstrate a highly contiguous, near-complete chromosome-scale resource suitable for comparative genomic and transcriptomic analyses.

Annotation of *B. juncea* identified a total of 15,159 coding genes and 20,046 isoforms (Supplementary Table 1: Genes (AGAT)). Mean gene length was 7,930 bp with a mean of 1.3 transcripts per gene, figures consistent with *P. dominula*^42^ (Supplementary Table 1: Genes (AGAT)). Annotation completeness assessed using the hymenoptera_odb10 BUSCO database reached 97.8% (Figure 2; Supplementary Table 1).

#### Mitochondrial genome

MitoHiFi assembled a single circularised mitochondrial genome of 21,580 bp containing 35 genes (**Supplementary Figure 1**). This mitogenome size and gene complement are consistent with those reported for other Vespidae, confirming accurate mitochondrial assembly and providing a complete reference sequence for population-level or phylogenetic studies of *B. juncea*. The raw sequence was deposited on Genbank (PZ371018).

### 2. Comparisons with other social wasps

#### Genome quality and gene content

Orthology-based phylogenetic inference using Orthofinder on extracted coding sequences showed clear grouping of branches in Vespidae subfamilies: Polistinae (species *B. juncea[this paper]* and *P. dominula*^42^) and Vespinae (genus *Dolichovespula*^83–85^, *Vespa*^43^ and *Vespula*^86,87^) (**Figure 2**; for samples see Supplementary table 1 [Comparison genomes]). *Ancistrocerus nigricornis (A. nigricornis)*^88^ and *L. flavolineata*^41^ (Eumeninae and Stenogastrinae subfamilies, respectively) appear as outgroups with regards to both Polistinae and Vespinae species, consistent with the literature^29^.

Total length of the other Vespidae assemblies ranged from 179 Mb to 370 Mb, and the GC content ranged from 30.79% to 41.62%, so the *B. juncea* assembly is well within the expected biological range for this family (Supplementary Table 1: Quast). Genome completeness was uniformly high across the 12 assemblies (BUSCO 94.8–98.4% complete; *B. juncea* 98.0%). The *B. juncea* annotation was among the most complete, recovering 97.8% of BUSCO genes from the predicted proteins (86.0–98.2% in the other species), with few duplicated BUSCOs (0.9%) (Figure 2; Supplementary Table 1). *B. juncea* had the lowest scaffold N50 (7.9 Mb, compared with 8.5–40.2 Mb), which reflects its larger number of smaller chromosomes rather than poorer contiguity (see *Chromosome number*, below).

*B. juncea* also had the largest gene set (15,159 genes, compared with 9,858–13,686 in the other species), approximately 2,000 genes more than reported in *P. dominula*^42^ (13,089 genes; Supplementary Table 1: Genes (AGAT)). It had few overlapping gene models (5.7%), whereas overlapping models were common in some Vespinae annotations (e.g. 53% in *V. germanica*). Much of this excess lies in a few *B. juncea*-specific gene families whose coding sequences largely overlap annotated transposable elements (see Gene expansions and contractions, below). Methodologically, differences in annotation pipelines, repeat masking stringency, and the depth of RNA-seq training data used across assemblies can each inflate or deflate gene counts.

#### Transposable elements and repeats

Using the Dfam transposable-element families for Hymenoptera, RepeatMasker masked 32.0% of the *B. juncea* genome, within the range of the other vespids (15.2–46.6%) and similar to *P. dominula* (34.8%; *L. flavolineata* 15.2%; **Figure 2**; **Supplementary Table 1: RepeatMasker**), although Dfam includes species-specific repeat libraries for all of the other species except *L. flavolineata*, so repeats are likely to be under-estimated in *B. juncea*. Classified transposable elements made up similar proportions of the genomes. DNA transposons spanned 7.2% of the *B. juncea* genome (7.3% in both *P. dominula* and *L. flavolineata*), with Tc1–IS630–Pogo the most common (2.7%); LTR retrotransposons spanned 3.4% (3.2% and 1.3%), mostly Gypsy/DIRS1 (2.3%); and LINEs spanned 2.1% (2.3% and 0.4%).

This library-based approach misses the largest repeat in the *B. juncea* genome. The assembly contains arrays of a satellite with a 296-bp repeat unit (consensus sequence: https://github.com/Sumner-lab/Belonogaster_genome_paper/blob/main/HiC_and_satellite/satellite_296bp_consensus.fa), present on all 39 chromosomes in 685 arrays of up to 2.8 Mb. This satellite is not in Dfam: RepeatMasker masked only 2.7% of it. The de novo estimate of repeat content from HiTE is 50.3% (Supplementary Table 1: HiTE). The satellite was not detected in any of the other 11 vespid assemblies. Omni-C reads cannot be mapped uniquely within such long arrays of near-identical repeats, so the arrays appear as pale bands in the contact map (Figure 1b): 87% of the satellite lies in regions with almost no Omni-C contacts. We hesitantly assume that these repeats represent the centromeres of the chromosomes, but more investigation is required.

#### Chromosome number

The Omni-C contact map revealed 39–43 well-defined interaction blocks along the diagonal, indicating strong intra-chromosomal contact and successful chromosome-scale scaffolding (**Figure 1b**). Orthology analysis identified 39 scaffolds containing at least five BUSCO genes, which we designate as pseudo-chromosomes (**Figure 2**). This chromosome count is markedly higher than the 22–26 chromosomes typical of other sequenced Vespidae, including 25 chromosomes in *Vespula* species^87^ and 22 in *P. dominula*^42^ (**Supplementary Table 1**: Comparison genomes), and it explains the lower scaffold N50 of the *B. juncea* assembly. The range of 39–43 blocks comes from the two largest scaffolds (HiC_scaffold_1 and 2; **Figure 1b**), each of which contains several contact blocks that are separated by satellite arrays (see Transposable elements and repeats, above) and interact only weakly with one another.

The elevated chromosome number in *B. juncea* is intriguing and warrants careful interpretation. Three non-mutually exclusive explanations exist: first, *B. juncea* may genuinely possess a higher karyotype, reflecting lineage-specific chromosomal fission events following divergence from other Polistinae. Such events are known in other Hymenoptera and need not be accompanied by changes in genome size, consistent with our observation that the *B. juncea* total assembly size falls within the range seen across Vespidae. Second, some of the 39 pseudo-chromosomes could represent incompletely scaffolded arms of larger chromosomes — a limitation that Hi-C-based assembly approaches can face when contact signal across centromeric or heterochromatic regions is sparse, as it is across the satellite arrays described above. Third, and most likely, both factors contribute. Direct cytological karyotyping of *B. juncea* would be valuable to resolve this question and is an important avenue for future work. Regardless, the syntenic and gene-content analyses presented below suggest that the 39 scaffolds are functionally coherent chromosomal units rather than fragmented artefacts.

#### Synteny

To explore the syntenic relationship between *B. juncea* and related species, we ran a nextflow pipeline (Eco-flow/synteny v4.0.1^89^) to compare the regions of the genome that correspond between *B. juncea* (39 orthologous chromosomes), *P. dominula*^42^ (22 chromosomes, only 21 orthologous), *L. flavolineata*^41^ (13 chromosomes) and a vespine wasp, *Vespula vulgaris*^90^ *(V. vulgaris;* 25 chromosomes) (**Figure 3**). *L. flavolineata*, being the most distantly related wasp, shows little synteny compared to *B. juncea* as would be expected (**Figure 3-top pair**)*. B. juncea* appears to have shorter stretches of syntenic genes compared to *P. dominula* than *P. dominula* does to *V. vulgaris* (**Figure 3**), despite the former pair being more closely related. This is due to the breaking up of chromosomes in *B. juncea*. *B. juncea* compared to *V. vulgaris* has 9,525 syntenic genes in 446 syntenic blocks, whereas *P. dominula* compared to *V. vulgaris* has 9,292 syntenic genes in 329 syntenic blocks. These comparisons suggest that *B. juncea* has undergone considerable genome evolution, since diverging from the common ancestor shared with the *Polistes* clade, breaking up previously longer stretches of synteny (Raw data in Zenodo: https://doi.org/10.5281/zenodo.20704168).

**Figure 3.**
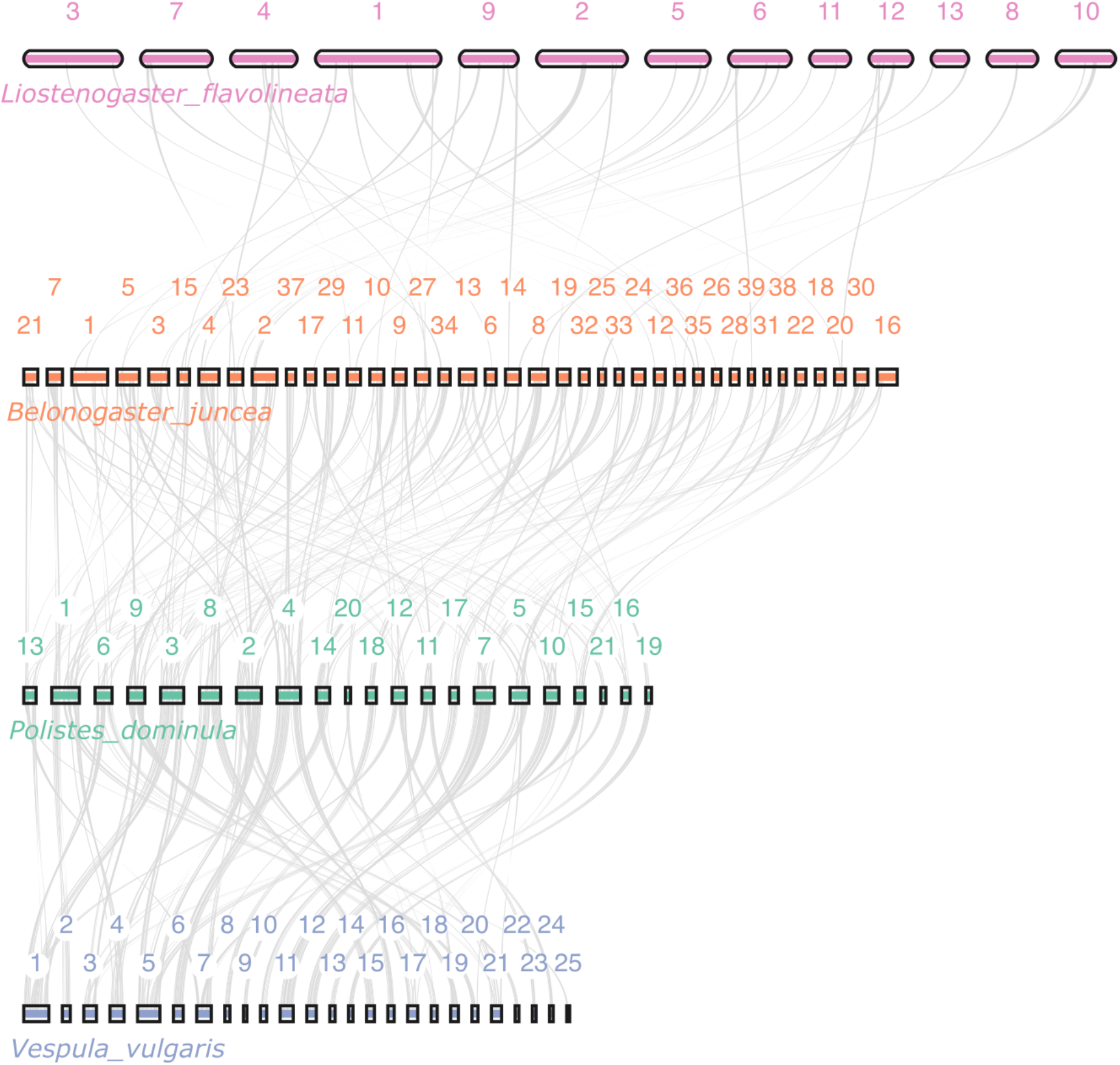
Syntenic relationships between *B. juncea*, *P. dominula*, *L. flavolineata* and *V. vulgaris*. For each species, the chromosomes are represented as horizontal bars, with the length relatively equivalent to their size. Grey bands stretching between pairs of chromosomes represent segments of DNA that are syntenic between species. Labels represent the chromosome names (as listed in the genome of each species), except *V. vulgaris* where a key to the chromosome name is found in Supp. Table 1: “Chromosome_names”.

#### Gene expansions and contractions

To explore the high gene count in *B. juncea*, we performed a gene expansion and contraction analysis with CAFE5^82^ using a pipeline Excon^79^ v2.5.1 (commit b809d7c). Many wasps in this clade have a number of significantly (*P*-value < 0.05) expanding and contracting orthogroups (OGs; **Figure 4a**; Supplementary Table 1: CAFE). *B. juncea* had 129 significantly expanded OGs, second only to *Vespa crabro* (140), but these expansions added far more genes than in any other species: 1,182, compared with 423 in *P. dominula* and 334 in *V. crabro*. *B. juncea* also had very few significant contractions (2, compared with 9–354 in the other species except *L. flavolineata*, whose branch cannot be interpreted due to being at the base of the tree; Figure 4a). About half of the gain (605 genes) lies in 32 *B. juncea*-specific families whose genes are short, often single-exon, rarely carry GO annotations and are rarely expressed in the brain. Their coding sequences largely overlap transposable elements annotated by RepeatMasker (median 89% of each gene’s coding sequence, mainly LINE, DNA and LTR elements and unclassified repeats, compared with 0% for single-copy orthologues; Methods), so much of this expansion reflects transposable-element-derived gene models rather than functional gene duplication. The results were robust to the definition of gene families: with hierarchical orthogroups from OrthoFinder v2.5.5^69^, *B. juncea* again showed by far the largest gain (+1,442 genes in 132 significantly expanded families, compared with at most 436 in the other species) and only two significant contractions. Full CAFE analysis files, including the OrthoFinder v2 analysis, are on Zenodo: https://doi.org/10.5281/zenodo.23373652.

**Figure 4.**
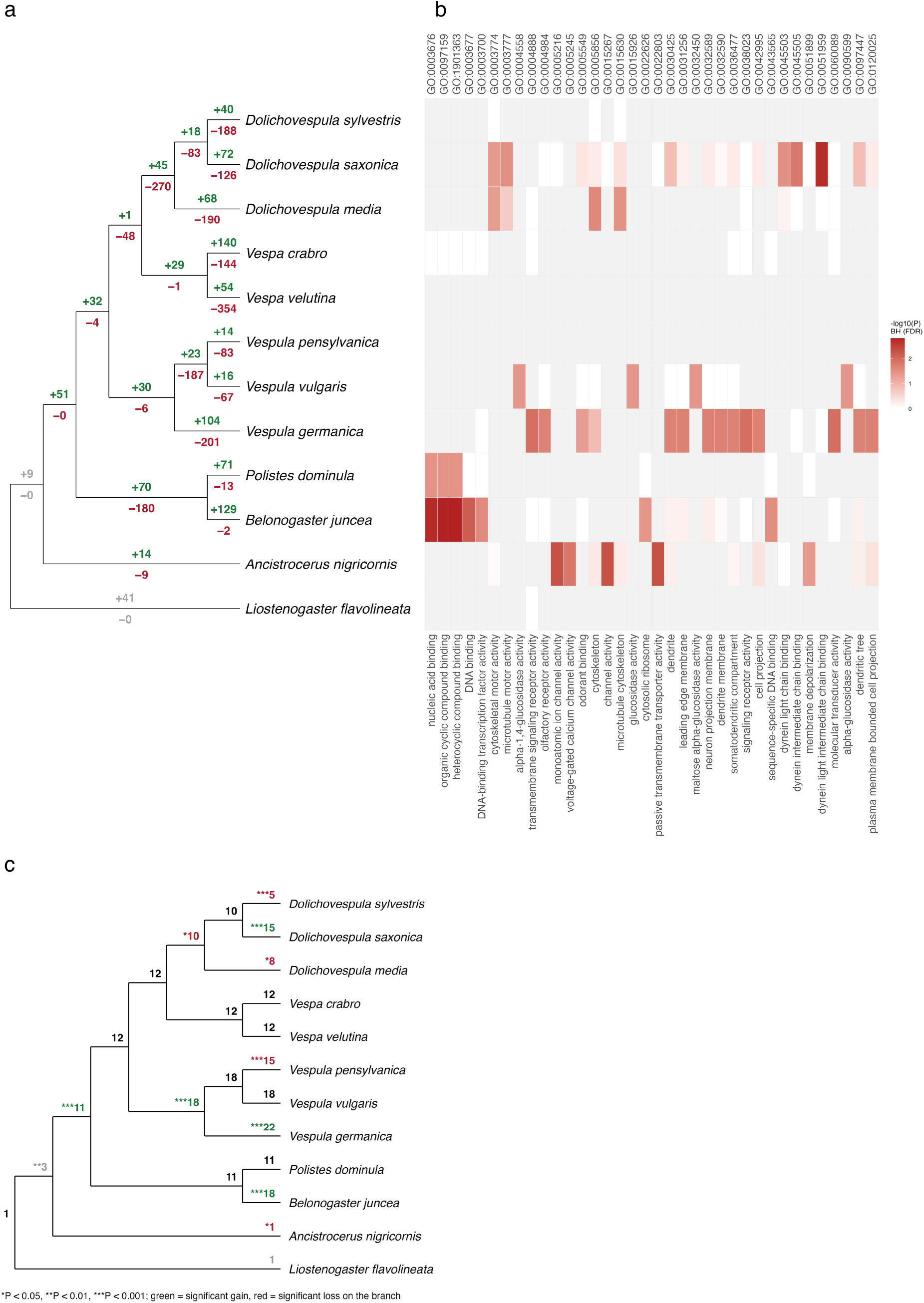
Gene family expansions and contractions. (a) OrthoFinder species tree, drawn with equal branch lengths, with the number of gene families (orthogroups) significantly expanded (green, +) or contracted (red, −) on each branch (CAFE5, branch P ≤ 0.05). The two branches either side of the root (grey) cannot be separated, because *L. flavolineata* is the only species on one side of the root, and are not interpreted. (b) GO terms enriched among the gene families expanded in each species (any increase), compared with all families in that species (topGO, classic Fisher’s exact test). Colour shows −log₁₀ of the Benjamini–Hochberg-corrected *P* (false discovery rate, corrected across all terms tested); terms with FDR < 0.05 in at least one species are shown; grey, not significant before correction. (c) Copy number of the largest olfactory receptor family (OG0000009) at each tip and ancestral node, as reconstructed by CAFE5. Coloured numbers mark a significant gain (green) or loss (red) on the branch leading to that node (\**P* < 0.05, \*\**P* < 0.01, \*\*\**P* < 0.001); grey, root branches not interpreted.

For GO enrichment we used all OGs that gained genes in each species, whether or not the gain was individually significant (e.g. 339 in *B. juncea*; Supplementary Table 1: CAFE). The terms enriched after correction for all terms tested (Benjamini–Hochberg false discovery rate, FDR < 0.05) differed between species rather than following phylogenetic relationships (Figure 4b). In *B. juncea*, expanded OGs were enriched for nucleic acid binding (GO:0003676; 43 OGs observed vs 21.8 expected; FDR = 0.0018), DNA binding (GO:0003677; FDR = 0.0071), DNA-binding transcription factor activity (GO:0003700; FDR = 0.028) and cytosolic ribosome (GO:0022626; FDR = 0.039). These came from small gains, mostly of one or two genes, in many conserved families, not from the *B. juncea*-specific families above. *P. dominula*, the other polistine, showed the same nucleic acid binding enrichment (FDR = 0.037). Expanded OGs in other species were enriched for ion channel activity (*A. nigricornis*), dynein binding and microtubule motor activity (*D. saxonica*), cytoskeleton (*D. media*), alpha-glucosidase activity (*V. vulgaris*) and signalling receptor and neuronal membrane terms (*V. germanica*). Among contracted OGs (Supplementary Figure 2), ribosomal terms were enriched in *V. pensylvanica*, nucleosome and chromatin terms in *V. germanica*, and cytosolic ribosome in *B. juncea* (FDR = 0.040). Results for all species are on Zenodo (https://doi.org/10.5281/zenodo.23373652; “cafe_go” folder), and lists of the gene families expanded and contracted in each species are in “cafe_plot/cafe_plotter/gene_family.tar.gz” (same record).

Odorant receptors are among the most dynamic gene families in social Hymenoptera^91^, including *Polistes* paper wasps^92^, so we also examined them directly. Olfactory GO terms were not enriched among *B. juncea* expansions overall, but the largest olfactory receptor family (OG0000009) was significantly expanded on the *B. juncea* branch (18 genes, compared with 11 in *P. dominula*; CAFE P = 1.0 × 10⁻^4^; Figure 4c). The family expanded at the base of the Polistinae and Vespinae (from 3 to 11 copies) and changed repeatedly thereafter, with independent gains in *Vespula* and *D. saxonica* and losses in other Vespinae (Figure 4c). Olfactory terms were enriched among expanded OGs in *V. germanica* (olfactory receptor activity, GO:0004984, FDR = 0.022; odorant binding, GO:0005549, FDR = 0.0499) and among contracted OGs in *V. pensylvanica* (FDR ≤ 0.012), consistent with frequent gains and losses of these genes across vespids. All 18 *B. juncea* copies lie in a 63-kb tandem array on chromosome 13, which accounts for the enrichment of olfactory receptor activity on that chromosome (see Functional enrichment of chromosomes, below).

#### Functional enrichment of chromosomes

Given the unusually high number of chromosomes in *B. juncea* supported by our HiC results (Figure 1b), we asked if distinct chromosomes have distinct functional specialisations. Using gene ontology (GO) analysis, we tested whether the genes on each chromosome were enriched for particular functions compared with all genes in the genome (Bonferroni-corrected P-value < 0.05; Figure 5). Counting all genes, 12 of the 39 chromosomes showed enrichment for specific functions (Zenodo: https://doi.org/10.5281/zenodo.23221104). For instance, scaffold 16 is enriched for mRNA splicing via the spliceosome (20.6% of GO:0000398, Bonferroni-corrected *P*-value: 1.33e-35), scaffold 2 for the small-subunit processome (52.9% of GO:0032040, Bonferroni-corrected P-value: 1.84e-24), scaffold 13 for olfactory receptor activity (65.4% of GO:0004984, Bonferroni-corrected P-value: 2.79e-16), and scaffold 24 for cuticle hydrocarbon biosynthesis (46.2% of GO:0006723, Bonferroni-corrected P-value: 9.75e-04). However, most of these enrichments reflected clusters of duplicated genes. When each orthogroup was counted only once per chromosome, only four chromosomes retained an enriched term, and only one strongly: scaffold 20, enriched for midgut development (27.6% of GO:0007494, Bonferroni-corrected P-value: 6.60e-05). Similar patterns were found all other vespid wasps in this study (samples: **Supplementary Table 1** [Comparison genomes]; result: https://doi.org/10.5281/zenodo.23221104), suggesting that the extra chromosomes of *B. juncea* do not necessarily signify functional sub-compartmentalisation. Comparisons with more closely-related wasps are needed to explore if any of the more recent chromosome fissions have led to some dichotomy.

**Figure 5.**
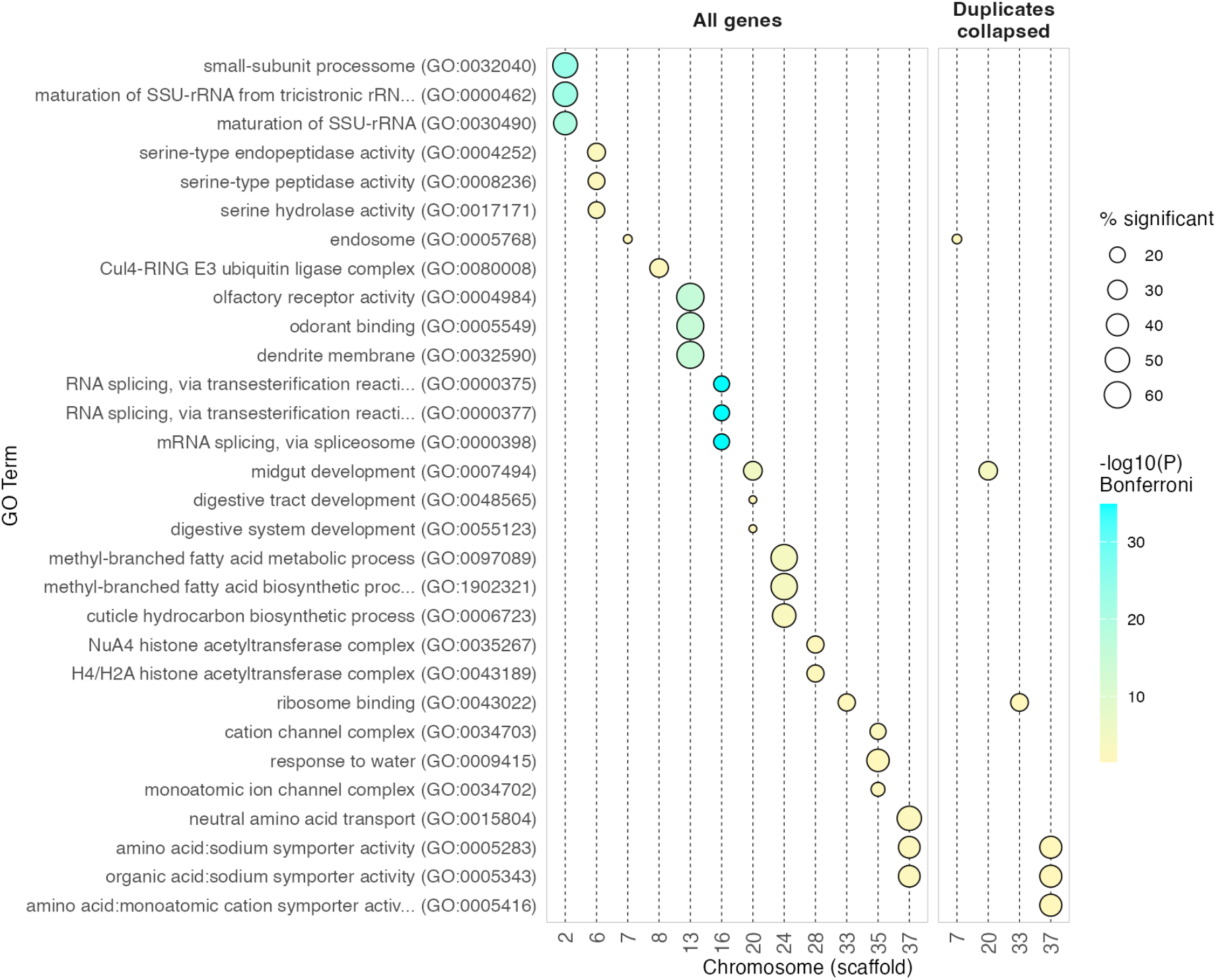
Gene ontology (GO) enrichment of *B. juncea* chromosomes. For each chromosome (scaffold), the top three GO terms enriched among its genes compared with all genes in the genome (one-sided Fisher’s exact test; Bonferroni-corrected *P* < 0.05 across all terms tested; terms with ≥ 10 annotated genes). Left: all genes counted. Right: each orthogroup counted once per chromosome, so clusters of duplicated genes contribute only once. Point size shows the percentage of the term’s genes (or orthogroups) located on that chromosome; colour shows −log10 of the Bonferroni-corrected *P*.

### 3. Caste-specific brain transcription and comparisons with other social wasps

#### Caste-specific gene expression

To investigate the molecular basis of reproductive division of labour, we quantified gene expression in 110 individual reproductives and non-reproductives across all three species, retaining 10,243 (67.6%) genes in *B. juncea [this paper]*, 10,934 (79.9%) in *L. flavolineata*^72^, and 9,254 (70.7%) in *P. dominula*^73^ after filtering for low expression (**Supplementary Table 2**). After dimension reduction via PCA, reproductives and nonreproductives in *B. juncea* and *L. flavolineata* exhibited relatively weak separation, with stronger clustering in *P. dominula* (**Figure 6**).

**Figure 6.**
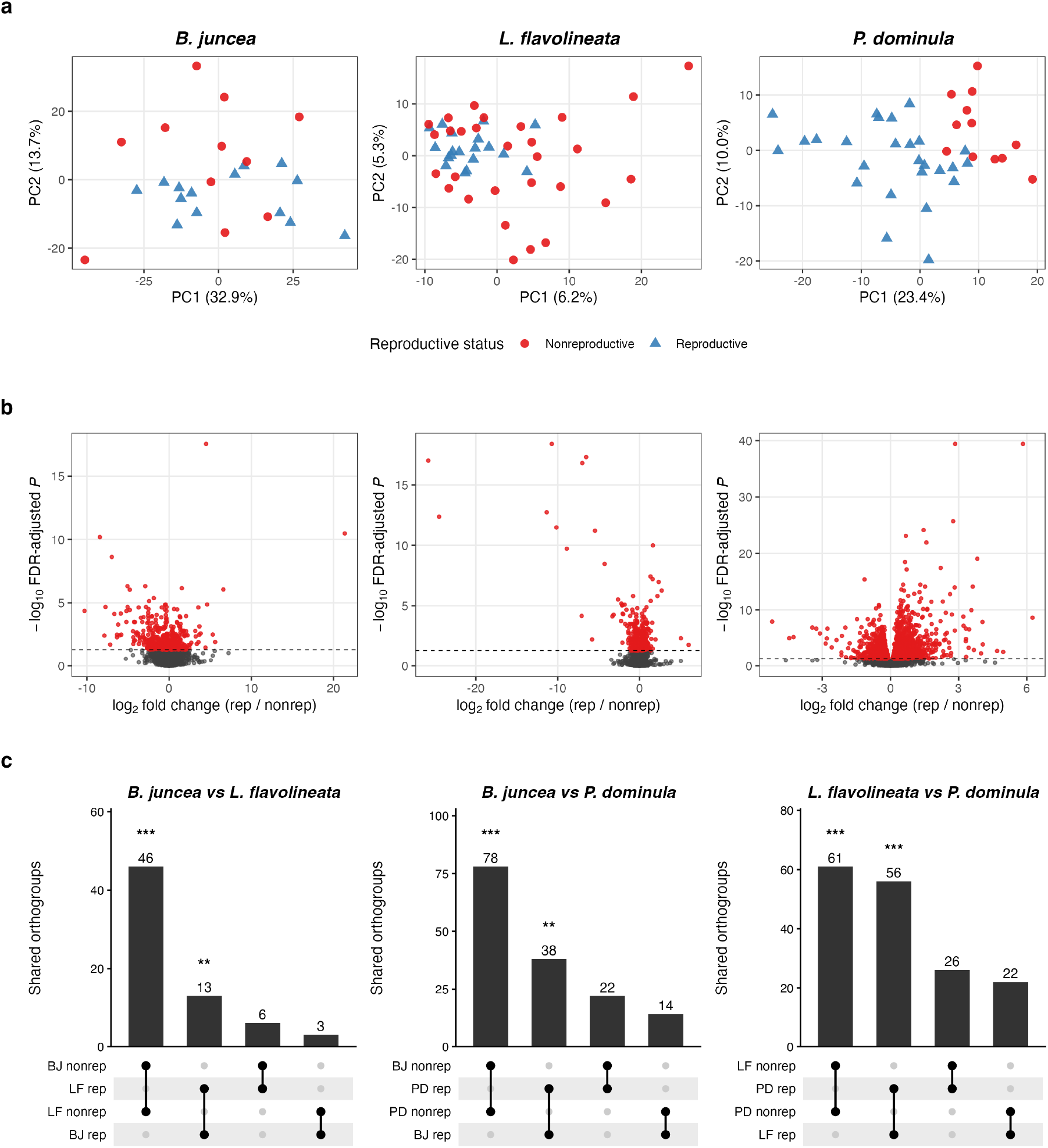
Differential expression between castes. (a) Principal component analysis of brain gene expression (variance-stabilised counts of the 500 most variable genes) in *B. juncea*, *L. flavolineata* and *P. dominula*; red circles, non-reproductives; blue triangles, reproductives. (b) Volcano plots of differential expression between reproductives and non-reproductives (DESeq2): log2 fold change (reproductive / non-reproductive) against −log10 FDR-adjusted P; red, genes with FDR < 0.05 (dashed line). (c) Numbers of orthogroups shared between pairs of species that contain genes upregulated in reproductives (rep) or non-reproductives (nonrep) in each species. Asterisks mark same-caste overlaps larger than expected by chance (one-sided hypergeometric test; *P < 0.05, **P < 0.01, ***P < 0.001). Full results and the shared orthogroups are in Supplementary Table 2 (Orthogroup_overlap_tests and Orthogroup_overlap_shared).

Next, we identified genes differentially expressed between reproductives and non-reproductives in each species at an FDR-corrected significance threshold of 0.05: 573 in *B. juncea* (232 upregulated in reproductives, hereafter queen-biased, and 341 upregulated in non-reproductives, hereafter worker-biased), 583 in *L. flavolineata* (276 and 307) and 2,416 in *P. dominula* (1,366 and 1,050; Figure 6; Supplementary Table 2). In *B. juncea*, all ten samples from Soa were reproductives, so caste is partly confounded with collection site; however, when the model was refitted on the 15 samples from Yaoundé alone, 98.8% of the 573 DEGs changed in the same direction (Spearman correlation of fold changes 0.97; Supplementary Table 2: Belonogaster_site_check). To compare species, we assigned DEGs to orthogroups and tested whether pairs of species shared more caste-biased orthogroups than expected by chance (Methods). Orthogroups containing worker-biased genes overlapped strongly between all three pairs of species (2.0–4.5 times more than expected; *B. juncea* and *L. flavolineata*, 46 shared orthogroups vs 10.1 expected, P = 1 × 10^⁻18^; *B. juncea* and *P. dominula*, 78 vs 35.7, P = 5 × 10⁻^12^; *L. flavolineata* and *P. dominula*, 61 vs 31.3, P = 1 × 10⁻^7^), and orthogroups containing queen-biased genes overlapped more weakly (1.5–2.6 times; P = 3 × 10⁻^5^–0.010). Orthogroups that were queen-biased in one species and worker-biased in another did not overlap more than expected (P ≥ 0.87). Thus, although the specific genes differ among species, there is some convergence in the transcriptional programmes associated with reproductive status, especially among non-reproductives.

Functional enrichment of differentially expressed gene sets identified significantly enriched gene ontology (GO) terms (FDR < 0.05) in each of the species (**Figure 7**; **Supplementary Table 3**), with *B. juncea* specifically having an enrichment of phototransduction-related terms among worker-biased genes (GO:0007602, FDR-corrected P = 8.1 × 10⁻^7^; of the 39 expressed genes annotated to this term in *B. juncea*, 13 were upregulated in non-reproductives, compared with 1.40 expected by chance). Although we interpret these results as preliminary given the small sampling sizes and collection of queens from different sites/times of day, this may reflect sensory differences in caste-specific roles, where visual processing demands may be especially acute for workers who conduct foraging^93^. While olfactory processes are known to be especially important in complex insect societies, which live in enclosed nests where visual cues are less useful and who use chemical cues in communication^94^; visual receptors may be especially important for wasps as they rely heavily on visual perception in their hunting and dominance behaviours^95^.

**Figure 7.**
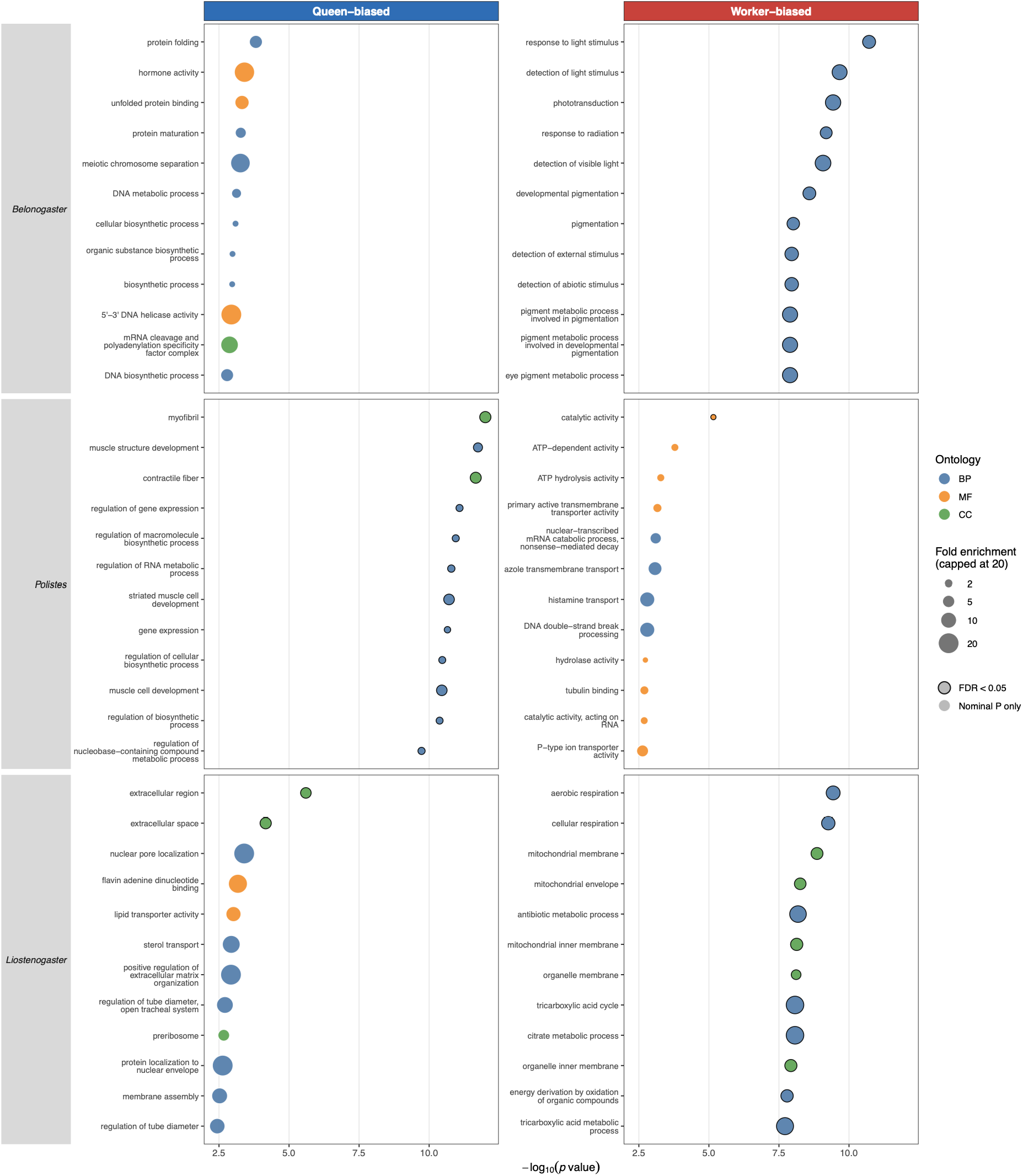
Gene Ontology (GO) terms enriched among caste-biased genes. For each species (rows), the 12 most significantly enriched GO terms (one-sided Fisher’s exact test, P < 0.01) among genes upregulated in reproductives (queen-biased; left) and in non-reproductives (worker-biased; right). The x-axis shows −log₁₀(*P*-values). Point colour denotes the GO domain (BP, biological process; MF, molecular function; CC, cellular component). Point size shows fold enrichment, i.e. the observed number of caste-biased genes annotated to the term divided by the number expected by chance (capped at 20). Black outlines mark terms that remain significant after Benjamini–Hochberg correction across all terms tested in that domain (FDR < 0.05). Enrichment was tested against a background of expressed genes. Full results are in Supplementary Table 3.

The GO terms identified were diverse across species and largely non-overlapping (**Supplementary Table 3**; **Supplementary Figure 3**). The exception was worker-biased genes in *B. juncea* and *L. flavolineata*, which shared 23 enriched terms (P < 0.01), mostly related to energy metabolism (e.g. the tricarboxylic acid cycle). This is more than expected by chance (4.2 expected; permutation test, BH-adjusted P = 0.02). No other pair of gene sets shared more terms than expected after correction, and no term was enriched in the same caste in all three species. The absence of a conserved enriched functional signature across all three species is consistent with some previous comparative studies of social insects^96^ suggesting that the molecular pathways underpinning caste are evolutionarily labile, with different lineages co-opting distinct gene regulatory networks to achieve broadly similar social outcomes^97^.

#### Conclusion

We present a high-quality chromosome-scale genome assembly and annotation for *B. juncea*, the first for this genus and the first for any aculeate wasp from the Global South. The assembly is among the most complete available for Vespidae, and provides a foundation for comparative analyses of genome evolution, gene regulation, and the molecular basis of sociality across the wasp phylogeny. The unexpectedly high chromosome number in *B. juncea* is a notable finding that merits cytological follow-up. Patterns of caste-associated gene expression - and their shared and contrasting features with other simple social wasp species - add to a growing body of evidence that the plastic nature of castes in simple societies are regulated by a shared ‘toolkit’ of conserved genes, alongside lineage and ecology-specific selection. Together, these resources and analyses position *B. juncea* as an important comparative model for the study of social evolution in Hymenoptera, and contribute to building capacity for exciting new research avenues for researchers in the Global South.

## Data availability

The *Belonogaster juncea* genome assembly and raw sequencing reads have been deposited at NCBI under BioProject PRJNA1357342 (GenBank assembly accession GCA_057814355.1; SRA run accessions SRR39278268 for PacBio HiFi and SRR41179205 for Omni-C). RNA-seq reads and processed expression matrices are available from NCBI GEO under accession GSE320383 (BioProject PRJNA1427691). Genome assemblies, gene annotations (V1 and V2; V2 used in this study) and protein sequences are available on Zenodo for *B. juncea* (https://doi.org/10.5281/zenodo.18892456), *P. dominula* (https://doi.org/10.5281/zenodo.23264930) and *L. flavolineata* (https://doi.org/10.5281/zenodo.23266517). The *V. crabro* Hi-C genome assembly and gene annotation used for comparisons are available on Zenodo (https://doi.org/10.5281/zenodo.23374842). All code used for the analyses is available at https://github.com/Sumner-lab/Belonogaster_genome_paper and archived on Zenodo (https://doi.org/10.5281/zenodo.23417166) under the MIT licence. The Juicer/3D-DNA contact map and assembly layouts used for Figure 1b, the Tandem Repeats Finder output and the consensus sequence of the 296-bp satellite are available on Zenodo (https://doi.org/10.5281/zenodo.23378133).

## Author contributions

SS, CW, conceived and designed the paper.

RT, MT, PK & PS sourced and collected samples.

SS, AT, SR, PK, CW acquired funding for the work.

CO did the dissections, ovary activation.

CW wrote the manuscript, with revisions from PK and SS.

CW, FD, BAT, EF, KG & RM performed data analysis.

AC performed data curation and uploaded genomes to NCBI.

All authors reviewed, edited and approved the final manuscript.

## Funding and acknowledgments

We thank the Cameroonian Ministry of Environment, Nature Protection, and Sustainable Development (MINEPDED, permit 00001/MINEPDED/CNA/NP-ABS/ABS-FP to PK), Ministry of Scientific Research and Innovation (MINRESI, permits 000042/MINRESI/B00/C00/C10/C13 and 000048/MINRESI/B00/C00/C10/C13 to PK) and the Ministry of Forests and Fauna (MINFOF, permit 0890PRBS/MINFOF/SETAT/SG/DFAP/SDVEF/SC/BJ to PK). This work was supported by the UKRI Natural Environment Research Council (NE/S011218/1 to SS, AT, SR and UKRI1263 to PK), US National Science Foundation (NE/S011218/1 to SS, AT, SR), ERC Horizon Europe (MSCA WaspClim 895220, to PK and ERC Starting Grant SOCIALEARTH, 101221668, to PK), and UKRI Biotechnology and Biological Sciences Research Council (BB/X018768/1 to SS, CW). Views and opinions expressed are those of the authors only and do not necessarily reflect those of the European Union or the European Research Council. Neither the European Union nor the granting authority can be held responsible for them. This work was supported by an award from BioFAIR to CW. BioFAIR is a joint Biotechnology and Biological Sciences Research Council (BBSRC) and Medical Research Council (MRC) project. It has received funding through the UKRI Infrastructure Fund and UKRI Digital Research Infrastructure Programme.

## Supplementary Figures

**Supplementary Figure 1:**
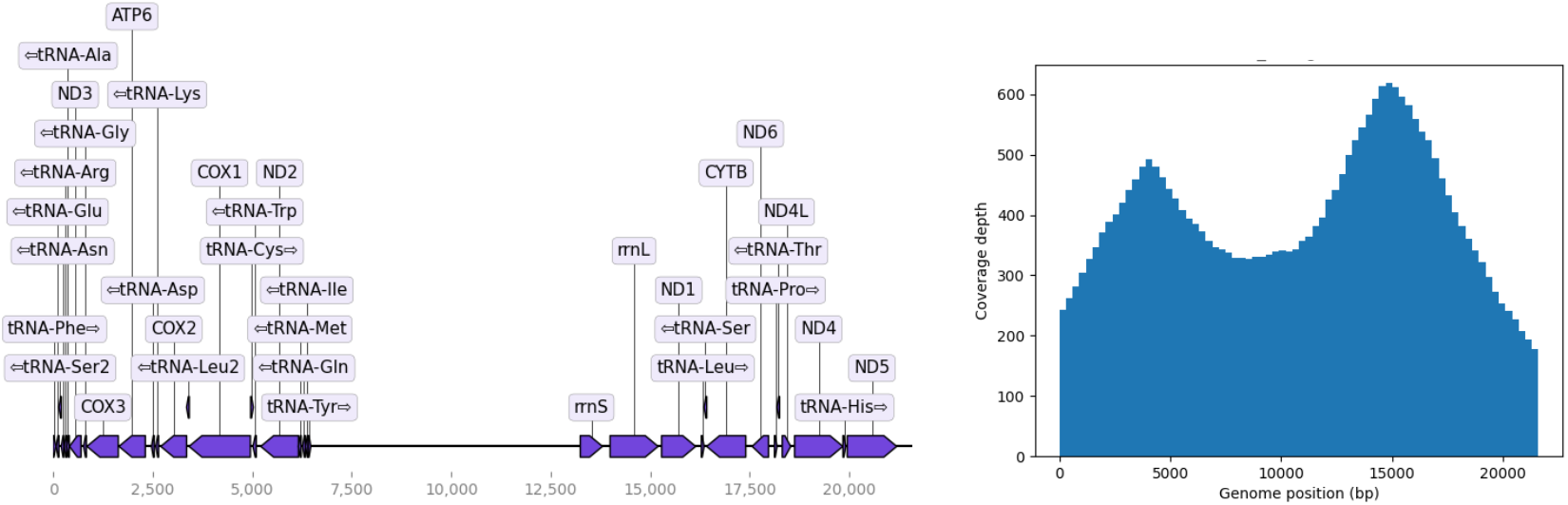
*B. juncea* mitochondrial genome assembly. The mitochondrion comprised 35 annotated genes, with a total genome size of 21,580 bp. Plotted using *MitoHiFi*.

**Supplementary Figure 2.**
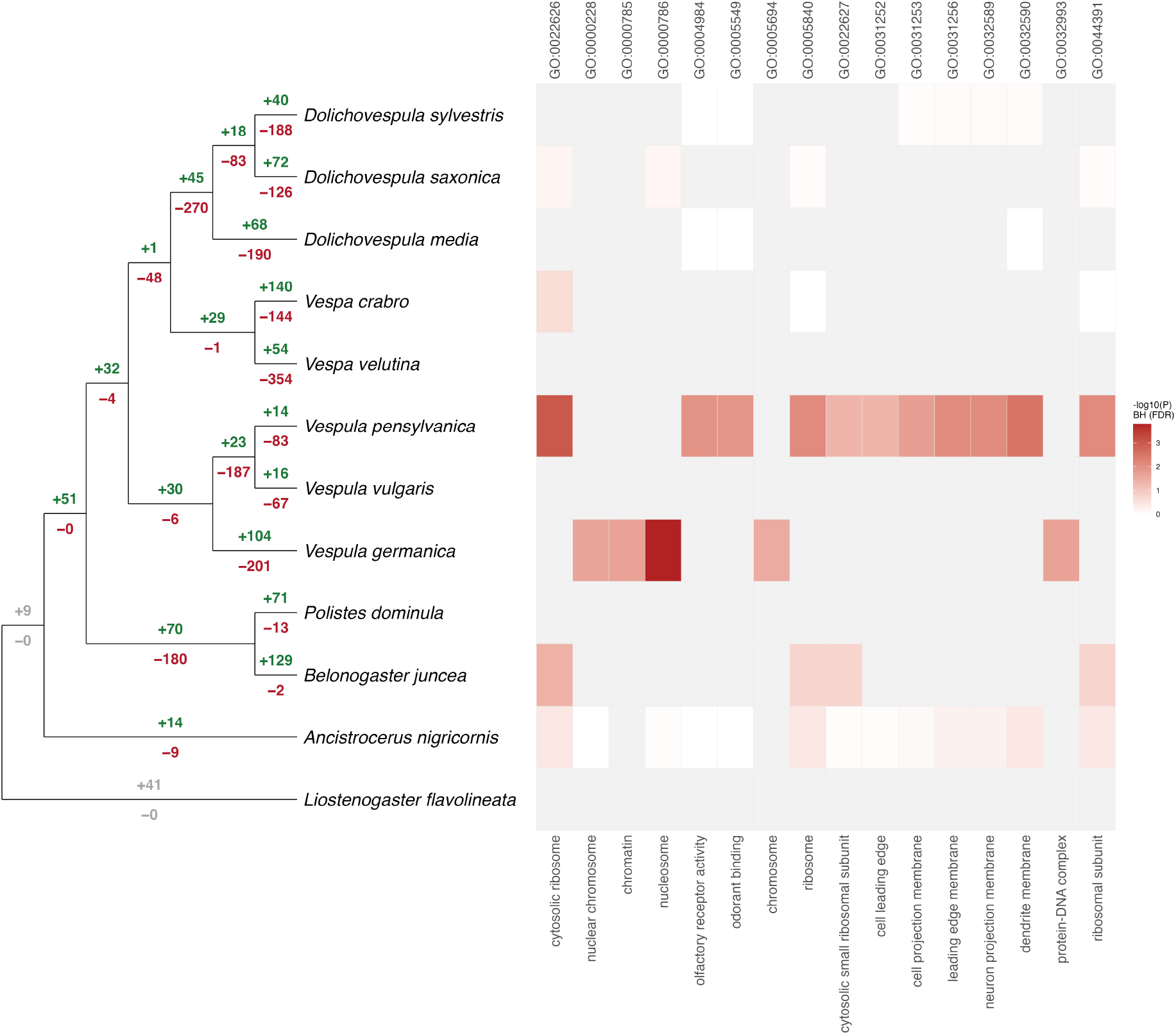
G**O enrichment of contracted gene families.** As Figure 4a and b, for gene families contracted in each species (any decrease). Terms with Benjamini– Hochberg false discovery rate < 0.05 in at least one species are shown.

**Supplementary Figure 3.**
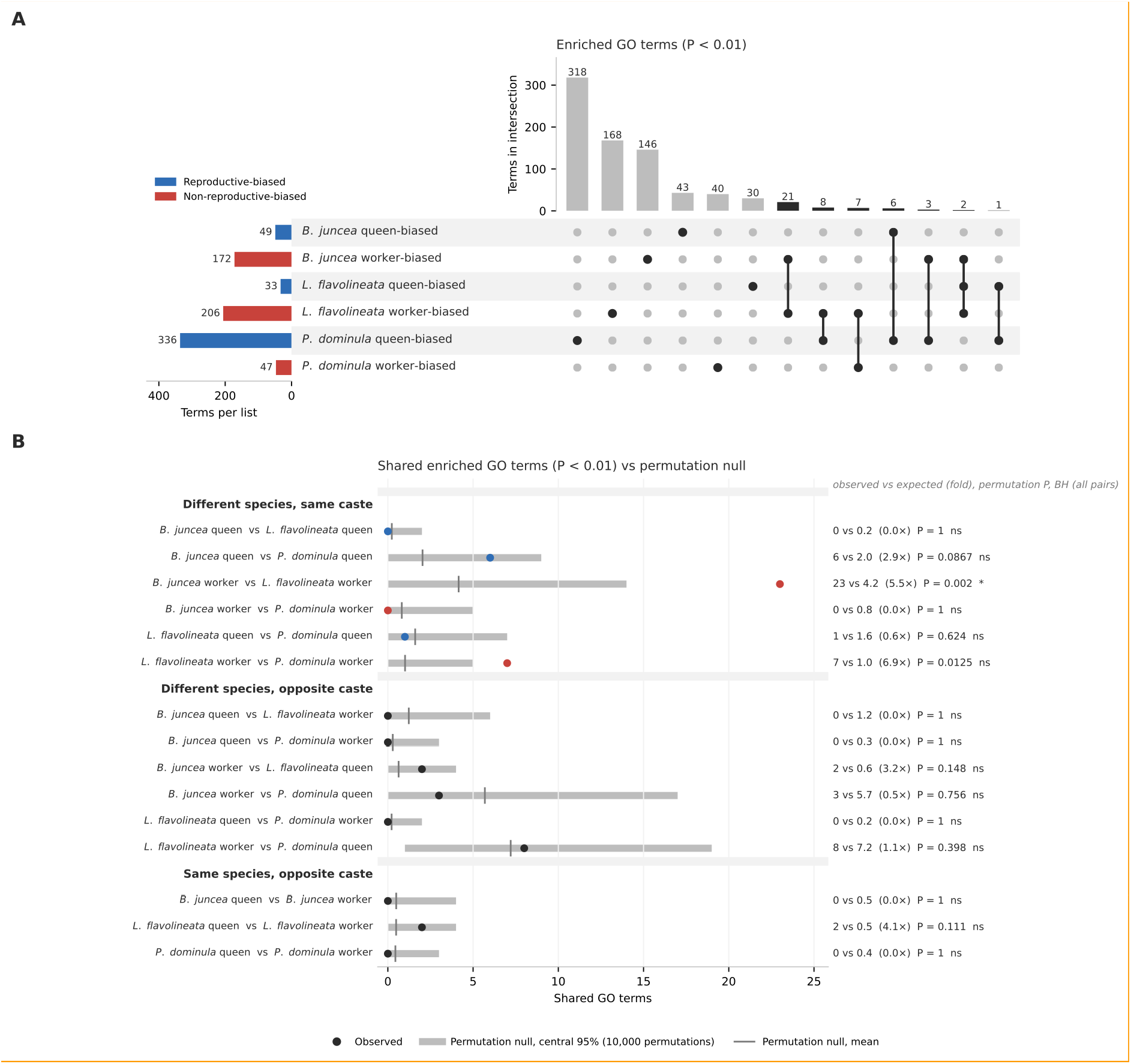
Overlap of enriched GO terms between caste-biased gene sets. (A) UpSet plot of GO terms enriched (P < 0.01) in each gene set (left bars; blue, queen-biased; red, worker-biased). Top bars count terms enriched in exactly the gene sets marked below. (B) Enriched terms shared by each pair of gene sets from different species (points) compared with chance expectation (grey bar, central 95%; line, mean), from 10,000 permutations of random gene sets (see Methods). Right: observed vs expected shared terms, fold enrichment and permutation P; *BH-adjusted P < 0.05. Counts in (B) include terms also enriched in other gene sets.

## Citations

1. Hunt, J. H. & Toth, A. L. Sociality in Wasps. in Comparative Social Evolution (eds Rubenstein, D. R. & Abbot, P.) 84–123 (Cambridge University Press, 2017). doi:10.1017/9781107338319.005.

2. Jandt, J. M., Tibbetts, E. A. & Toth, A. L. *Polistes paper* wasps: a model genus for the study of social dominance hierarchies. Insectes Sociaux 61, 11–27 (2014).

3. Bell, E. & Sumner, S. Ecology and Social Organisation of Wasps. in Encyclopedia of Life Sciences (Wiley, 2013). doi:10.1002/9780470015902.a0023597.

4. Patalano, S. et al. Molecular signatures of plastic phenotypes in two eusocial insect species with simple societies. Proc. Natl. Acad. Sci. 112, 13970–13975 (2015).

5. Standage, D. S. et al. Genome, transcriptome and methylome sequencing of a primitively eusocial wasp reveal a greatly reduced DNA methylation system in a social insect. Mol. Ecol. 25, 1769–1784 (2016).

6. Miller, S. E. et al. Evolutionary dynamics of recent selection on cognitive abilities. Proc. Natl. Acad. Sci. 117, 3045–3052 (2020).

7. Miller, S. E., Legan, A. W., Uy, F. M. K. & Sheehan, M. J. Highly Contiguous Genome Assemblies of the Guinea Paper Wasp *Polistes exclamans* and *Mischocyttarus mexicanus*. Genome Biol. Evol. 14, evac110 (2022).

8. Gilligan, J. et al. Rapid and cost-effective production of high-quality insect genome sequences. Preprint at 10.1101/2024.10.06.616902 (2024).

9. Linck, E. B. & Cadena, C. D. A Latitudinal Gradient of Reference Genomes. Mol. Ecol. e17551 (2024) doi:10.1111/mec.17551.

10. Tindo, M., D’Agostino, P., Francescato, E., Dejean, A. & Turillazzi, S. Associative colony foundation in the tropical wasp *Belonogaster juncea juncea* (Vespidae, Polistinae). Insectes Sociaux 44, 365–377 (1997).

11. Tindo, M. & Dejean, A. Dominance hierarchy in colonies of *Belonogaster juncea juncea* (Vespidae, Polistinae): Insectes Sociaux 47, 158–163 (2000).

12. Tindo, M., Mony, R. & Dejean, A. Colony Development and Serial Polygyny in the Primitively Eusocial Wasp *Belonogaster juncea juncea* (Vespidae: Polistinae). J. Insect Behav. 15, 243–252 (2002).

13. Tindo, M., Kenne, M. & Dejean, A. Advantages of multiple foundress colonies in *Belonogaster juncea juncea* L.: greater survival and increased productivity. Ecol. Entomol. 33, 293–297 (2008).

14. Richards, O. W. A revision of the genus *Belonogaster* de Saussure. Bull. Br. Mus. Nat. Hist. Entomol. 44, 31–114 (1982).

15. Roubaud, É. Recherches biologiques sur les guêpes solitaries et sociales d’Afrique. Ann. Sci. Nat. Zool. (1916).

16. Keeping, M. G. Colony Foundation and Nestmate Recognition in the Social Wasp, *Belonogaster petiolata*. Ethology 85, 1–12 (1990).

17. Keeping, MalcolmG. Social organization and division of labour in colonies of the polistine wasp, *Belonogaster petiolata*. Behav. Ecol. Sociobiol. 31, (1992).

18. Keeping, M. G. Social behavior and brood decline in reproductive-phase colonies of *Belonogaster petiolata* (Degeer) (Hymenoptera: Vespidae). J. Insect Behav. 10, 265–278 (1997).

19. Kelstrup, H. C., Hartfelder, K., Esterhuizen, N. & Wossler, T. C. Juvenile hormone titers, ovarian status and epicuticular hydrocarbons in gynes and workers of the paper wasp *Belonogaster longitarsus*. J. Insect Physiol. 98, 83–92 (2017).

20. du Buysson, R. Monographie des Vespides du Genre Belonogaster. 78, 199–270.

21. Wheeler, W. M. A Study of Some Ant Larvæ, with a Consideration of the Origin and Meaning of the Social Habit among Insects. Proc. Am. Philos. Soc. 57, 293–343 (1918).

22. Roubaud, É. Évolution de l’instinct chez les Vespides: aperçus biologiques sur les Guêpes sociales d’Afrique du genre *Belonogaster* Sauss. Comptes Rendus Hebd. Séances Académie Sci. 151, 553–556 (1910).

23. Thomson, J. A. Secrets of Animal Life. (Henry Holt and Company, New York, 1919).

24. Marino Piccioli, M. T. & Pardi, L. Studies on the Biology of *Belonogaster* (Hymenoptera, Vespidae): 3. The Nest of Belonogaster griseus (Fab.). Monit. Zool. Ital. Suppl. 10, 179–228 (1978).

25. Marino Piccioli, M. T. & Pardi, L. Studi sulla biologia di *Belonogaster* (Hymenoptera, Vespidae): 1. Sull’etogramma di Belonogaster griseus (Fab.). Monit. Zool. Ital. Suppl. 3, 197–225 (1970).

26. Pardi, L. & Marino Piccioli, M. T. Studi sulla biologia di *Belonogaster* (Hymenoptera, Vespidae): 2. Differenziamento castale incipiente in B. griseus (Fab.): Pubblicazioni del Centro di Studio sulla Faunistica e la Ecologia dell’Africa Orientale del C.N.R.: XLIX. Monit. Zool. Ital. Suppl. 3, 235–265 (1970).

27. Pardi, L. & Marino Piccioli, M. T. Studies on the Biology of *Belonogaster* (Hymenoptera, Vespidae): 4. On Caste Differences in Belonogaster griseus (Fab.) and the Position of This Genus Among Social Wasps: Pubblicazioni del Centro di Studio per la Faunistica ed Ecologia Tropicali del C.N.R.: CXCVI. Monit. Zool. Ital. Suppl. 14, 131–146 (1981).

28. Keeping, M. G. Reproductive and worker castes in the primitively eusocial wasp *Belonogaster petiolata* (DeGeer) (Hymenoptera: Vespidae): evidence for pre-imaginal differentiation. J. Insect Physiol. 48, 867–879 (2002).

29. Piekarski, P. K., Carpenter, J. M., Lemmon, A. R., Moriarty Lemmon, E. & Sharanowski, B. J. Phylogenomic Evidence Overturns Current Conceptions of Social Evolution in Wasps (Vespidae). Mol. Biol. Evol. 35, 2097–2109 (2018).

30. Cheng, H., Concepcion, G. T., Feng, X., Zhang, H. & Li, H. Haplotype-resolved de novo assembly using phased assembly graphs with hifiasm. Nat. Methods 18, 170–175 (2021).

31. Ruan, J. & Li, H. Fast and accurate long-read assembly with wtdbg2. Nat. Methods 17, 155–158 (2020).

32. Laetsch, D. R. & Blaxter, M. L. BlobTools: Interrogation of genome assemblies. F1000Research 6, 1287 (2017).

33. De Coster, W., D’Hert, S., Schultz, D. T., Cruts, M. & Van Broeckhoven, C. NanoPack: visualizing and processing long-read sequencing data. Bioinformatics 34, 2666–2669 (2018).

34. Li, H. Aligning sequence reads, clone sequences and assembly contigs with BWA-MEM. Preprint at 10.48550/ARXIV.1303.3997 (2013).

35. Broad Institute. Picard toolkit. (2019).

36. Durand, N. C. et al. Juicer Provides a One-Click System for Analyzing Loop-Resolution Hi-C Experiments. Cell Syst. 3, 95–98 (2016).

37. Dudchenko, O. et al. De novo assembly of the *Aedes aegypti* genome using Hi-C yields chromosome-length scaffolds. Science 356, 92–95 (2017).

38. Durand, N. C. et al. Juicebox Provides a Visualization System for Hi-C Contact Maps with Unlimited Zoom. Cell Syst. 3, 99–101 (2016).

39. Wyatt, C. D. R. PacBio HiFi sequencing of Belonogaster juncea (isolate Bel_junc_adF). NCBI Sequence Read Archive (2026).

40. Wyatt, C. D. R. et al. nf-core/genomeqc: a best-practice pipeline for comparing genome and assembly quality. Preprint at 10.64898/2026.08.20.745971 (2026).

41. Wyatt, C. D. R., Geist, K., Favreau, E. & Masonbrink, R. Liostenogaster flavolineata genome assembly and annotation (PRJNA1415745). NCBI GenBank (2026).

42. Wyatt, C. D. R., et al. Data: Genome assembly, gene prediction and associated proteins of P. dominula. Zenodo 10.5281/zenodo.23264930 (2026).

43. Wyatt, C. Data: Hi-C genome assembly and gene prediction of Vespa crabro (Sumner lab). Zenodo 10.5281/ZENODO.23374842 (2026).

44. Gurevich, A., Saveliev, V., Vyahhi, N. & Tesler, G. QUAST: quality assessment tool for genome assemblies. Bioinformatics 29, 1072–1075 (2013).

45. Simão, F. A., Waterhouse, R. M., Ioannidis, P., Kriventseva, E. V. & Zdobnov, E. M. BUSCO: assessing genome assembly and annotation completeness with single-copy orthologs. Bioinformatics 31, 3210–3212 (2015).

46. Smit, A. F. A.; Hubley, R.; Green, P. RepeatMasker Open-4.0. http://www.repeatmasker.org (2013).

47. Storer, J., Hubley, R., Rosen, J., Wheeler, T. J. & Smit, A. F. The Dfam community resource of transposable element families, sequence models, and genome annotations. Mob. DNA 12, 2 (2021).

48. Steinegger, M. & Söding, J. MMseqs2 enables sensitive protein sequence searching for the analysis of massive data sets. Nat. Biotechnol. 35, 1026–1028 (2017).

49. Benson, G. Tandem repeats finder: a program to analyze DNA sequences. Nucleic Acids Res. 27, 573–580 (1999).

50. Flynn, J. M. et al. RepeatModeler2 for automated genomic discovery of transposable element families. Proc. Natl. Acad. Sci. 117, 9451–9457 (2020).

51. Dobin, A. et al. STAR: ultrafast universal RNA-seq aligner. Bioinformatics 29, 15–21 (2013).

52. Hoff, K. J. & Stanke, M. WebAUGUSTUS--a web service for training AUGUSTUS and predicting genes in eukaryotes. Nucleic Acids Res. 41, W123–W128 (2013).

53. Hoff, K. J., Lomsadze, A., Borodovsky, M. & Stanke, M. Whole-Genome Annotation with BRAKER. in Gene Prediction (ed. Kollmar, M.) vol. 1962 65–95 (Springer New York, New York, NY, 2019).

54. Lomsadze, A., Burns, P. D. & Borodovsky, M. Integration of mapped RNA-Seq reads into automatic training of eukaryotic gene finding algorithm. Nucleic Acids Res. 42, e119– e119 (2014).

55. Gremme, G., Brendel, V., Sparks, M. E. & Kurtz, S. Engineering a software tool for gene structure prediction in higher organisms. Inf. Softw. Technol. 47, 965–978 (2005).

56. Buchfink, B., Reuter, K. & Drost, H.-G. Sensitive protein alignments at tree-of-life scale using DIAMOND. Nat. Methods 18, 366–368 (2021).

57. Hu, K. et al. HiTE: a fast and accurate dynamic boundary adjustment approach for full-length transposable element detection and annotation. Nat. Commun. 15, 5573 (2024).

58. Quinlan, A. R. & Hall, I. M. BEDTools: a flexible suite of utilities for comparing genomic features. Bioinformatics 26, 841–842 (2010).

59. Bolger, A. M., Lohse, M. & Usadel, B. Trimmomatic: a flexible trimmer for Illumina sequence data. Bioinformatics 30, 2114–2120 (2014).

60. Ewels, P., Magnusson, M., Lundin, S. & Käller, M. MultiQC: summarize analysis results for multiple tools and samples in a single report. Bioinformatics 32, 3047–3048 (2016).

61. Kim, D., Paggi, J. M., Park, C., Bennett, C. & Salzberg, S. L. Graph-based genome alignment and genotyping with HISAT2 and HISAT-genotype. Nat. Biotechnol. 37, 907– 915 (2019).

62. Danecek, P. et al. Twelve years of SAMtools and BCFtools. GigaScience 10, giab008 (2021).

63. Stanke, M., Steinkamp, R., Waack, S. & Morgenstern, B. AUGUSTUS: a web server for gene finding in eukaryotes. Nucleic Acids Res. 32, W309–W312 (2004).

64. Gabriel, L. et al. BRAKER3: Fully automated genome annotation using RNA-seq and protein evidence with GeneMark-ETP, AUGUSTUS and TSEBRA. Preprint at 10.1101/2023.06.10.544449 (2023).

65. Brůna, T., Lomsadze, A. & Borodovsky, M. GeneMark-ETP significantly improves the accuracy of automatic annotation of large eukaryotic genomes. Genome Res. 34, 757– 768 (2024).

66. Gabriel, L., Hoff, K. J., Brůna, T., Borodovsky, M. & Stanke, M. TSEBRA: transcript selector for BRAKER. BMC Bioinformatics 22, 566 (2021).

67. Jacques Dainat et al. AGAT: Another Gff Analysis Toolkit to handle annotations in any GTF/GFF format. NBISweden/AGAT: AGAT v1.6.1. Zenodo 10.5281/ZENODO.18230424 (2026).

68. Pertea, G. & Pertea, M. GFF Utilities: GffRead and GffCompare. F1000Research 9, 304 (2020).

69. Emms, D. M. & Kelly, S. OrthoFinder: phylogenetic orthology inference for comparative genomics. Genome Biol. 20, 238 (2019).

70. Uliano-Silva, M. et al. MitoHiFi: a python pipeline for mitochondrial genome assembly from PacBio high fidelity reads. BMC Bioinformatics 24, 288 (2023).

71. Allio, R. et al. MitoFinder: Efficient automated large-scale extraction of mitogenomic data in target enrichment phylogenomics. Mol. Ecol. Resour. 20, 892–905 (2020).

72. Taylor, B. A. et al. Molecular signatures of alternative reproductive strategies in a facultatively social hover wasp. Mol. Ecol. 33, e17217 (2024).

73. Taylor, B. A., Cini, A., Wyatt, C. D. R., Reuter, M. & Sumner, S. The molecular basis of socially mediated phenotypic plasticity in a eusocial paper wasp. Nat. Commun. 12, 775 (2021).

74. Harshil Patel et al. nf-core/rnaseq: nf-core/rnaseq v3.21.0 - Mercury Macaw. Zenodo 10.5281/ZENODO.1400710 (2025).

75. Di Tommaso, P. et al. Nextflow enables reproducible computational workflows. Nat. Biotechnol. 35, 316–319 (2017).

76. Patro, R., Duggal, G., Love, M. I., Irizarry, R. A. & Kingsford, C. Salmon provides fast and bias-aware quantification of transcript expression. Nat. Methods 14, 417–419 (2017).

77. Love, M. I., Huber, W. & Anders, S. Moderated estimation of fold change and dispersion for RNA-seq data with DESeq2. Genome Biol. 15, 550 (2014).

78. Benjamini, Y. & Hochberg, Y. Controlling the False Discovery Rate: A Practical and Powerful Approach to Multiple Testing. J. R. Stat. Soc. Ser. B Methodol. 57, 289–300 (1995).

79. Wyatt, C. D. R. et al. EXCON: a scalable, reproducible Nextflow pipeline for CAFE-based gene EXpansion and CONtraction analysis. 2026.09.25.754512 Preprint at 10.64898/2026.09.25.754512 (2026).

80. Cantalapiedra, C. P., Hernández-Plaza, A., Letunic, I., Bork, P. & Huerta-Cepas, J. eggNOG-mapper v2: Functional Annotation, Orthology Assignments, and Domain Prediction at the Metagenomic Scale. Mol. Biol. Evol. 38, 5825–5829 (2021).

81. Alexa, A., Rahnenführer, J. & Lengauer, T. Improved scoring of functional groups from gene expression data by decorrelating GO graph structure. Bioinformatics 22, 1600– 1607 (2006).

82. Mendes, F. K., Vanderpool, D., Fulton, B. & Hahn, M. W. CAFE 5 models variation in evolutionary rates among gene families. Bioinformatics 36, 5516–5518 (2021).

83. Falk, S. et al. The genome sequence of the tree wasp, *Dolichovespula sylvestris* Scopoli, 1763. Wellcome Open Res. 7, 113 (2022).

84. Wang, H. et al. Next-Generation Sequencing of Four Mitochondrial Genomes of *Dolichovespula* (Hymenoptera: Vespidae) with a Phylogenetic Analysis and Divergence Time Estimation of Vespidae. Animals 12, 3004 (2022).

85. Crowley, L. M. et al. The genome sequence of the Median Wasp, *Dolichovespula media* (Retzius, 1783). Wellcome Open Res. 8, 492 (2023).

86. Harrop, T. W. R. et al. High-Quality Assemblies for Three Invasive Social Wasps from the *Vespula* Genus. G3 GenesGenomesGenetics 10, 3479–3488 (2020).

87. Crowley, L. M. et al. The genome sequence of the German wasp, *Vespula germanica* (Fabricius, 1793). Wellcome Open Res. 7, 60 (2022).

88. Crowley, L. M. et al. The genome sequence of the Early Mason-wasp, *Ancistrocerus nigricornis* (Curtis, 1826). Wellcome Open Res. 8, 461 (2023).

89. Wyatt, C. A Nextflow pipeline to conduct a gene wise synteny analysis. Preprint at 10.48546/WORKFLOWHUB.WORKFLOW.2190.1 (2026).

90. Crowley, L. et al. The genome sequence of the common wasp, *Vespula vulgaris* (Linnaeus, 1758). Wellcome Open Res. 6, 232 (2021).

91. McKenzie, S. K. & Kronauer, D. J. C. The genomic architecture and molecular evolution of ant odorant receptors. Genome Res. 28, 1757–1765 (2018).

92. Legan, A. W., Jernigan, C. M., Miller, S. E., Fuchs, M. F. & Sheehan, M. J. Expansion and Accelerated Evolution of 9-Exon Odorant Receptors in *Polistes* Paper Wasps. Mol. Biol. Evol. 38, 3832–3846 (2021).

93. O’Donnell, S. et al. A test of neuroecological predictions using paperwasp caste differences in brain structure (Hymenoptera: Vespidae). Behav. Ecol. Sociobiol. 68, 529–536 (2014).

94. Pellen, Y. et al. Adaptive evolution of odorant receptors is associated with elaborations of social organization in ants. Mol. Biol. Evol. 43, msag103 (2026).

95. D’adamo, P. & Lozada, M. The importance of location and visual cues during foraging in the German wasp *(Vespula germanica* F.) (Hymenoptera: Vespidae). N. Z. J. Zool. 30, 171–174 (2003).

96. Wyatt, C. D. R. et al. Social complexity, life-history and lineage influence the molecular basis of castes in vespid wasps. Nat. Commun. 14, 1046 (2023).

97. Favreau, E. et al. Co-expression Gene Networks and Machine-learning Algorithms Unveil a Core Genetic Toolkit for Reproductive Division of Labour in Rudimentary Insect Societies. Genome Biol. Evol. 15, evac174 (2023).

